# Small Molecules Targeting the Disordered Transactivation Domain of the Androgen Receptor Induce the Formation of Collapsed Helical States

**DOI:** 10.1101/2021.12.23.474012

**Authors:** Jiaqi Zhu, Xavier Salvatella, Paul Robustelli

## Abstract

Castration-resistant prostate cancer (CRPC) is a lethal condition suffered by ∼35% of prostate cancer patients who become resistant to existing FDA-approved drugs. Small molecules that target the intrinsically disordered N-terminal domain of the androgen receptor (AR-NTD) have shown promise in circumventing CPRC drug-resistance. A prodrug of one such compound, EPI-002, entered human trials in 2015 but was discontinued after phase I due to poor potency. The compound EPI-7170 was subsequently found to have improved potency, and a related compound entered human trials in 2020. NMR measurements have localized the strongest effects of these compounds to a transiently helical region of the disordered AR-NTD but no detailed structural or mechanistic rationale exists to explain their affinity to this region or the comparative potency of EPI-7170. Here, we utilize all-atom molecular dynamics simulations to elucidate the binding mechanisms of the small molecules EPI-002 and EPI-7170 to the disordered AR-NTD. We observe that both compounds induce the formation of collapsed helical states in the Tau-5 transactivation domain and that these bound states consist of heterogenous ensembles of interconverting binding modes. We find that EPI-7170 has a higher affinity to Tau-5 than EPI-002 and that the EPI-7170 bound ensemble contains a substantially higher population of collapsed helical states than the bound ensemble of EPI-002. We identify a network of interactions in the EPI-7170 bound ensemble that stabilize collapsed helical conformations. Our results provide atomically detailed binding mechanisms for EPI compounds consistent with NMR experiments that will prove useful for drug discovery for CRPC.

**Summary:** Intrinsically disordered proteins (IDPs), which do not fold into a well-defined three-dimensional structure under physiological conditions, are implicated in many human diseases. Such proteins are difficult to characterize at an atomic level and are extremely challenging drug targets. Small molecules that target a disordered domain of the androgen receptor have entered human trials for the treatment of castration-resistant prostate cancer, but no structural or mechanistic rationale exists to explain their inhibition mechanisms or relative potencies. Here, we utilize molecular dynamics computer simulations to elucidate atomically detailed binding mechanisms of these compounds and understand their inhibition mechanisms. Our results suggest strategies for developing more potent androgen receptor inhibitors and general strategies for IDP drug design.

## Introduction

Intrinsically disordered proteins (IDPs) that lack a fixed three-dimensional structure under physiological conditions, represent ∼40% of the human proteome, have crucial functional roles in a variety of biological pathways and biomolecular assemblies and are implicated in many human diseases^1-8^. IDPs represent an enormous pool of potential drug targets that are currently inaccessible to conventional structure-based drug design methods utilized for folded proteins^9-15^. In conventional drug-design campaigns small molecules are designed to optimize the strength of intermolecular interactions with well-defined binding sites. The structures of drug binding sites in folded proteins are often known in their apo and bound forms from high-resolution crystal structures. In cases where bound structures of inhibitors are known, it is often straightforward to identify pharmacophores to optimize with small molecule modifications. A number of computational tools exist to predict the affinities of small molecules that bind structured binding sites in folded proteins, with alchemical free-energy perturbations calculations growing in popularity and offering increasingly accurate predictions relative to more high-throughput docking and virtual screening approaches^16-17^.

Unlike folded proteins, IDPs populate a conformational ensemble of rapidly interconverting structures in solution^18^. The conformations of IDPs can sample a large number of topological arrangements, and structures in IDP conformational ensembles can have little-to-no structural similarity to one another. Many IDPs possess sequence elements with elevated secondary structure populations relative to random-coil distributions^18,19^. This confers a limited amount of local order to IDP conformational ensembles, but the number of thermally accessible orientations of sidechain and backbone pharmacophores can remain combinatorially large. As such, it is generally not possible to represent even small sequence segments of IDPs by a single dominant conformation or a small number of substantially populated conformations. IDPs are therefore not suitable targets for conventional structure-based drug design methods that require the existence of an ordered binding site and the general “druggability” of IDP sequences remains uncertain^13^.

If it becomes possible to rationally target IDPs with small molecule drugs the druggable proteome will be dramatically expanded and therapeutic interventions may become accessible for currently untreatable diseases associated with aberrant biological interactions of IDPs such as those mediated by biomolecular condensate formation and protein misfolding^10,15,20-23^. Several small molecules have been discovered that interact with specific IDP sequences and inhibit their interactions^24-31^, and many of these compounds show clear biological activity^24-27^ Biophysical measurements, particularly small but reproducible NMR chemical shift perturbations observed in ligand titrations, indicate that these IDPs remain highly dynamic while interacting with these compounds^24-26,30,31^. These observations suggests that the sequence specificity of these ligands is conferred through a network of transient interactions, and that these bound states consist of heterogenous ensembles of interconverting binding modes.

A number of all-atom molecular dynamics (MD) simulation studies provide support for dynamic binding mechanisms between IDPs and small molecule ligands^30,32-36^. In several studies, ligands are observed to populate a broad distribution of binding modes that confer little-to-no detectable ordering in IDP binding sites relative to their apo forms^32-34^. In a recent joint MD and NMR study a series of small molecule ligands were found to bind specifically to the C-terminal region of α- synuclein with differing affinities without significantly altering the conformational ensembles of the apo and bound states^32^ This study proposed that the specificity and affinity of these ligands is conferred through a so-called *dynamic shuttling* mechanism where a ligand rarely forms multiple specific intermolecular interactions simultaneously and instead transitions among networks of spatially proximal interactions. In this mechanism, differences in the affinity and specificity of ligands are attributed to the complementarity of the orientations of protein and ligand pharmacophores in a dynamic IDP ensemble without evoking the notion of an ordered binding site. In some cases, biophysical experiments and molecular simulation data suggest that the conformational ensembles of IDPs undergo an entropic expansion upon ligand binding, where interactions with a small molecule can increase the number of conformations significantly populated by an IDP^27,28,36^. In other cases, biophysical experiments and simulation results suggest that small molecule ligands can cause a population-shift among existing IDP conformations^29^, drive the compaction of monomeric disordered state proteins upon binding^30,35^ or drive the formation of soluble oligomeric states^31^. The variety of binding mechanisms observed suggest that small molecules affect the conformational ensembles of an IDPs in a system dependent manner^10,11^.

While a number of small molecules have been found to bind and inhibit IDPs in vitro and in vivo one of the most pharmaceutically promising IDP:drug interactions is the inhibition of the disordered N-terminal transactivation domain (NTD) of the androgen receptor (AR) by a series of compounds with a common Bisphenol A scaffold^37-42^. AR is a large multidomain transcription factor that contains folded ligand-binding and DNA-binding domains in addition to the 558-residue intrinsically disordered NTD^24^. Transcriptionally active AR drives the growth of most prostate cancers^14^. In ∼65% of cases prostate cancer patients can be cured by surgery or radiotherapy. In the remaining ∼35% of patients, the progression of the disease can initially be treated with a number of FDA approved drugs that target the folded AR ligand binding-domain by competitively inhibiting the binding site of transcription activating androgens. After a period of 2-3 years however, these patients inevitably become refractory to pharmacological interventions and develop lethal castration-resistant prostate cancer (CRPC). In CRPC, cancer cells acquire mutations and express splice variants that enable the activation of AR at low levels of circulating androgens and in the presence of ligand-binding domain inhibitors. One of the most common CRPC resistance adaption is the expression of constitutively active AR splice variants that lack the ligand-binding domain entirely^43^, and therefore render all current FDA-approved drugs ineffective.

The discovery of small molecules that target the disordered AR-NTD provides a promising therapeutic strategy for CRPC, as a number of these compounds have been shown to inhibit the constitutively active splice variants of AR that lack the ligand-binding domain^14^. A series of AR-NTD inhibitors were discovered from screening libraries of natural products and were shown to have efficacy in CRPC mouse models^37-42^. Among these compounds EPI-002, a bisphenol A derivative later named Ralaniten, was identified as a promising AR-NTD inhibitor and a prodrug of EPI-002 (EPI-506; Ralaniten Acetate) was selected for phase I clinical trials in 2015^44^, becoming the first small-molecule known to directly bind an intrinsically disordered protein to be tested in humans^14^. Human trials were however discontinued after phase I due to insufficient potency and poor metabolic properties. The compound EPI-7170, a second-generation AR-NTD inhibitor, was subsequently shown to have dramatically improved potency and metabolic properties relative to EPI-002^40-42^, and an additional second-generation AR-NTD inhibitor with an undisclosed chemical structure (EPI-7386) entered phase I clinical trials in March 2020^45,46^.

Nuclear magnetic resonance (NMR) spectroscopy has localized the strongest interactions of EPI-002 to the transactivation unit 5 domain (Tau-5; residues A350-G448) of the AR-NTD^24^. The Tau-5 domain contains three regions with transiently populated helices termed R1, R2 and R3^24,47,48^. These domains were found to have the largest NMR chemical shift perturbations (CSPs) in EPI-002 titrations^24^. The atomic details of the molecular mechanisms by which EPI-002 and other EPI compounds interact with the Tau-5 are, however, not well understood and no structural or mechanistic rationale exists to explain their affinity to the AR-NTD or their relative efficacies in CRPC treatment. In the absence of such mechanistic understanding, it is not currently possible to rationally design new compounds with improved potencies or to utilize similar inhibition mechanisms to target other intrinsically disordered domains.

In this investigation, we utilize all-atom explicit solvent MD simulations with the state-of-the-art a99SB-*disp* force field^49^ to study the binding mechanisms of EPI-002 and EPI-7170 to the transiently helical Tau-5 region of the disordered AR-NTD. We observe that EPI-002 and EPI-7170 both induce the formation of compact helical molten-globule-like states in Tau-5. These bound states remain dynamic and sample a heterogenous ensemble of binding modes. We find that EPI-7170 has a 2.5-fold higher affinity to Tau-5 than EPI-002 and that the EPI-7170 bound ensemble is substantially more helical relative to the bound ensemble of EPI-002. We identify a network of interactions in the EPI-7170 bound ensemble that more effectively stabilize these collapsed helical conformations. Our results suggest that EPI compounds inhibit the activity of AR by inducing the partial folding of molecular recognition elements in Tau-5 domain into compact helical states and preventing interactions between AR and the transcriptional machinery required for elevated AR transactivation in CRPC patients. The atomically detailed binding mechanisms described here suggest strategies for developing more potent AR inhibitors for the treatment of CRPC.

## Results

### Molecular Dynamics Simulations of the Disordered Tau-5_R2_R3_ Region of the Androgen Receptor N-terminal Transactivation Domain are consistent with NMR experiments

Previously reported NMR chemical shift perturbations (CSPs) indicate that EPI-002 interacts non-covalently with a subset of three partially helical regions, termed R1, R2 and R3, in the transactivation unit 5 (Tau-5) domain (residues 350-448) of the disordered AR-NTD^24^. Circular Dichroism experiments and NMR secondary chemical shifts suggest that R1, R2 and R3 are approximately 10%, 30%, and 10% helical respectively when Tau-5 is in its apo form in solution^24,47,48,50^. The 31-residue R1 region (residues S341-G371) is separated from the 24-residue R2 region (residues L391-G414) by a 19-residue proline-rich linker region that appears to predominantly populate an extended polyproline II conformation based on sequence composition and secondary NMR chemical shifts^24,50^. The R2 region is separated from the R3 region (residues S426-G446) by a glycine rich linker (^415^AGAAGPGSGSP^425^) that is predicted to have no residual secondary structure propensity based on sequence and secondary chemical shifts (Fig. 1) ^24,50^.

**Figure 1.**
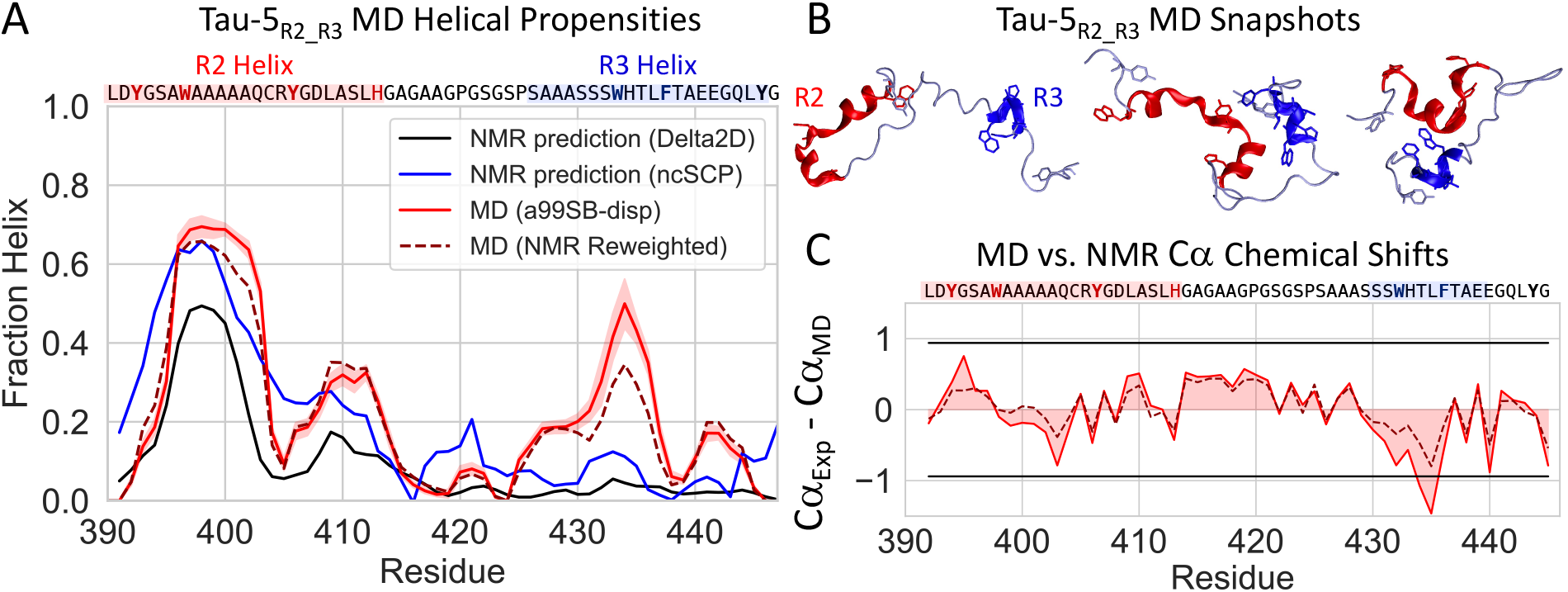
A) Comparison of the helical propensity observed in the 300K replica of a 74µs explicit solvent REST2 MD simulation of Tau-5_R2_R3_ in its apo form (red), the helical propensity of obtained after maximum-entropy reweighting of the MD ensemble using NMR Cα chemical shifts as restraints (discontinuous line, dark red) and helical propensity predictions derived from experimental NMR chemical shifts (blue and black).. Shaded regions indicate statistical error estimates from blocking. B) Illustrative snapshots from an MD simulation of the Tau-5_R2_R3_. The R2 and R3 regions of Tau-5_R2_R3_ are colored red and blue respectively. C) Comparison of experimental NMR Cα chemical shifts with shifts calculated from MD simulations using SPARTA+ before (red) and after maximum-entropy reweighting (discontinuous line, dark red). Black lines indicate the average Cα chemical shift prediction error of SPARTA+ on its training database of folded protein structures.

The relatively small magnitude of the backbone amide CSPs measured in Tau-5 upon EPI-002 binding suggest that this interaction does not induce rigid folded structures in these regions, and that the Tau-5 region remains highly dynamic while interacting with this compound^24^. Experiments were also performed to determine if the titration of peptides containing isolated R1, R2 and R3 domains produced CSPs or line broadening effects on the NMR resonances of EPI-001 (a racemic mixture containing EPI-002 along with three additional stereoisomers)^24^. No interactions were detected between EPI-001 and these truncated partially helical constructs, suggesting that multiple partially helical regions are required for the interaction between EPI-002 and Tau-5. These data suggest that EPI-002 may be binding at the interfaces of these transiently helical domains and stabilizing long-range tertiary contacts found in the apo Tau-5 ensemble.

The largest backbone amide NMR CSPs observed in EPI-002:Tau-5 titrations are found in the R2 and R3 regions. Specifically, the largest individual CSPs were found in the most helical residues of R2 (^400^AAQ^403^) and the ^433^WHTLF^437^ segment of R3. The ^433^WHTLF^437^ segment has been found to be critical for the function AR and to fold into a helical conformation upon binding the RAP74 domain of the general transcription regulator TFIIF^47,48,51,52^. The disruption of this interaction causes AR to lose is transcriptional activity^37,38,52^. Based on these observations and the sequence composition of Tau-5, we decided to focus our investigation on a 56-residue Tau-5 fragment (residues L391-G446) containing the R2 and R3 helices, which we refer to as Tau-5_R2_R3._ Given the extended nature of the proline-rich linker region between R1 and R2, and the small magnitude of CSPs in the R1 region upon EPI-002 binding^24^, we hypothesize that transient long-range contacts between R1 and R2 and R1 and R3 are less likely to be important for EPI-002 and EPI-7170 binding than shorter range contacts between R2 and R3. Simulating the smaller Tau-5_R2_R3_ fragment enables us to focus our investigation on the region of Tau-5 that showed the most pronounced CSPs in EPI-002 titrations and obtain better statistics on a more tractable sampling problem.

We ran an unbiased all-atom explicit solvent MD simulation of Tau-5_R2_R3_ with the a99SB-*disp* protein force field and a99SB-*disp* water model^49^ using the replica exchange with solute tempering (REST2) enhanced sampling algorithm^53^ (See Methods). Simulations with the a99SB-*disp* force field have been found to provide excellent agreement with the secondary structure propensities and the dimensions of IDPs. The apo Tau-5_R2_R3_ REST2 MD simulation was run with 16 replicas, spanning temperatures from 300K-500K, with all protein atoms selected for solute tempering. The simulation was run for 4.6µs/replica, for an aggregate simulation time of 74µs. The convergence of the simulation was assessed by computing statistical errors by a blocking analysis^55,56^ (See Methods) and comparing secondary structure propensities, intramolecular contact maps and free energy surfaces as a function of the radius of gyration (R_g_) and the α−helical order parameter Sα^54^ for each temperature rung in the REST2 simulation and for each independent demultiplexed replica (See Methods, Fig. S1-S6, Supporting Information: “Convergence Analyses”).

The average helical propensity observed in the REST2 MD simulation of apo Tau-5_R2_R3_ and illustrative snapshots from the 300K replica are shown in Fig. 1. The intramolecular contact map of Tau-5_R2_R3_ reveals relatively highly populated (∼60-80% occupancy) contacts between the partially helical ^396^AWAAAAAQCRY^406^ residues in R2 and the ^432^SWHTLF^437^ residues in R3 (Fig. S1). Statistical error estimates of the simulated properties of the apo Tau-5_R2_R3_ ensemble simulation were calculated using a blocking analysis following Flyvbjerg and Peterson^55,56^(See Methods). Using this approach, we calculated error estimates for secondary structure propensities (Average Helical Content=23.3 ± 0.6%), radius of gyration (R_g_=12.5 ± 0.1Å) and the helical order parameter Sα_54_ (Sα=6.5 ± 0.1). Sα reports on the similarity of each 6-residue fragment in the protein to an ideal helical conformation, with each 6-residue fragment with a small RMSD (<0.5 Å) from an ideal helical conformation contributing a value of ∼1 to the total Sα value (See Methods). The value of *S*α for a protein structure can therefore be interpreted as proxy for the number of 6-residue fragments closely resembling an ideal helical conformation. Together our convergence analyses (Fig. S1-S6) and statistical error estimates suggest that our apo Tau-5_R2_R3_ REST2 MD simulation is relatively well converged. A subset of conformations observed in the 300K replica of our apo Tau-5_R2_R3_ MD simulation are shown in Movie S1.

We validated the accuracy of our MD simulation of Tau-5_R2_R3_ by calculating NMR chemical shifts with SPARTA+^57^ and comparing them to previously reported experimental values^24^ (Table 1, Fig. 1). The overall agreement between calculated and experimental backbone chemical shifts is excellent and is on-par with the most accurate chemical shift predictions observed in force field benchmarks of simulations of IDPs^49,58^. The predicted Cα shifts are within the SPARTA+ Cα shift prediction error for nearly all residues (Fig. 1C), with the exception of small deviations in residues ^434^WH^435^. In addition to a direct comparison with NMR chemical shifts, we also observed that the simulated helical propensity of residues ^431^SSWHTLF^437^ in the R3 region are somewhat overestimated relative to helical propensities derived from NMR chemical shifts using the algorithms Delta2D and ncSCP ^59,60^ (Fig. 1A).

**Table 1.** Agreement between calculated and experimental NMR chemical shifts from the 300K replica of a 74µs unbiased REST2 MD simulation of Tau-5_R2_R3_ employing the a99SB-*disp* force field and from a maximum-entropy reweighted ensemble derived using Cα NMR chemical shifts as restraints. Chemical shifts were calculated using SPARTA+^57^.

| | Cα | HN | N | C' | Cβ |
| --- | --- | --- | --- | --- | --- |
| Unbiased Tau-5<sub>R2_R3</sub> MD Ensemble |  |  |  |  |  |
| RMSD (ppm) | 0.47 | 0.23 | 1.06 | 0.45 | 0.37 |
| Correlation | 0.991 | -0.558 | 0.954 | 0.915 | 1.000 |
| Cα-reweighted Maximum Entropy Tau-5<sub>R2_R3</sub> Ensemble | | | | | |
| RMSD (ppm) | 0.29 | 0.21 | 0.88 | 0.38 | 0.33 |
| Correlation | 0.997 | -0.312 | 0.968 | 0.934 | 1.000 |

To rigorously quantify the error in the simulated helical propensity directly against experimental data, we utilized the maximum-entropy reweighting algorithm of Cesari et al.^61,62^ (See Methods) to reweight our unbiased 300K ensemble using Cα chemical shifts as restraints (Table 1, Fig. 1). We note that while only Cα chemical shift predictions were restrained in the reweighting procedure, we observed improvements in the prediction accuracy of the remaining backbone shifts in the reweighted ensemble, suggesting that the resulting ensemble is not overfit to the Cα chemical shift data. We found that optimal agreement with experimental shifts was obtained by reducing the average helical propensity of residues ^431^SSWHTLF^437^ from 34% to 25%, suggesting an overstabilization of helical conformations in this region by 0.26kcal/mol (or ∼0.04 kcal/mol per residue in this 7-residue segment) in the unbiased trajectory. We note that the Cα-reweighted maximum entropy ensemble still possesses more helical content in the R3 region than is predicted by the NMR chemical shift based algorithms Delta2D and ncSCP (Fig. 1A). Considering that both chemical shift predictions from SPARTA+^57^ and NMR chemical shift based secondary structure prediction algorithms^57,59,60^ are subject to prediction errors, we consider the relatively modest deviation from experimental chemical shifts and predicted secondary structure propensities to be acceptable. Importantly, we note that while an unbiased Tau-5_R2_R3_ simulation run with the a99SB-*disp* force field somewhat overestimate the stability of the R3 helix, the simulated helical conformations in these region are only marginally stable, and we therefore expect to resolve increases or decreases in the stability of these conformations in the presence of ligands.

### EPI-7170 has a higher affinity to Tau-5_R2_R3_ than EPI-002

We utilized the same REST2 MD simulation protocol used for apo simulations of Tau-5_R2_R3_ to perform unbiased simulations of Tau-5_R2_R3_ in the presence of EPI-002 and EPI-7170 (See Methods). EPI-002 and EPI-7170 were parameterized with the GAFF1 forcefield^63^ Simulations run with GAFF1 and a99SB-*disp* force fields have previously been shown to provide excellent agreement with residue specific IDP ligand binding propensities based on comparisons to NMR CSPs^30,32^. A REST2 simulation of EPI-002 and Tau-5_R2_R3_ was run for 4.0µs/replica, for an aggregate simulation time of 64µs, and a REST2 simulation of EPI-7170 and Tau-5_R2_R3_ was run for 4.5µs/replica, for an aggregate simulation time of 72µs. The simulated properties of Tau-5_R2_R3_ observed in these simulations are compared to the apo Tau-5_R2_R3_ simulations in Table 2. Simulation convergence analyses are reported in Fig. S7-S21.

**Table 2.** Simulated values and statistical error estimates for properties of Tau-5_R2_R3_ in its apo form and when bound to EPI-002 and EPI-7170. Sα is an α−helical order parameter that is a proxy for the number of 6-residue helical fragments present in a protein conformation. Collapsed “helical globule” states are defined as Tau-5_R2_R3_ conformations with values of Sα>6.0 and R_g_<1.3nm. ΔG_globule_ is the free energy of formation of the helical globule state at 300K. Values for apo Tau-5_R2_R3_ were calculated from a simulation of Tau-5_R2_R3_ performed in the absence of ligands. Values for the EPI-002:Tau-5_R2_R3_ and EPI-7170:Tau-5_R2_R3_ bound ensembles were calculated using only bound frames in simulations in the presence of ligands. Error estimates were computed using a blocking analysis^55,56^.

| | $R_g$<br>(nm) | Fraction<br>Helix | $S\alpha$ | Bound<br>Fraction | Ligand $K_D$<br>(mM) | Helical Globule<br>Population | $\Delta G_{\text{globule}}$<br>(kcal/mol) |
| --- | --- | --- | --- | --- | --- | --- | --- |
| Apo Tau-5<sub>R2_R3</sub> | $1.25 \pm .01$ | $23.3 \pm 0.6\%$ | $6.5 \pm 0.1$ | - | - | $40.4 \pm 5.2\%$ | $+0.23 \pm 0.13$ |
| EPI-002: Tau-5<sub>R2_R3</sub><br>Bound Ensemble | $1.26 \pm 0.3$ | $25.4 \pm 1.3\%$ | $7.3 \pm 0.4$ | $43 \pm 2\%$ | $5.24 \pm 0.43$ | $48.5 \pm 6.5\%$ | $+0.04 \pm 0.17$ |
| EPI-7170: Tau-5<sub>R2_R3</sub><br>Bound Ensemble | $1.24 \pm .02$ | $32.8 \pm 0.5\%$ | $9.6 \pm 0.2$ | $67 \pm 2\%$ | $1.92 \pm 0.15$ | $61.1 \pm 4.5\%$ | $-0.27 \pm 0.11$ |

To calculate a simulated K_D_ value for each compound, we define the bound population (P_b_) of each ligand as the fraction of frames with at least one intermolecular contact between a and Tau-5_R2_R3_, where intermolecular contacts are defined as occurring in frames where at least one ligand heavy (non-hydrogen) atom is within 6.0Å of a protein heavy atom. This cutoff was selected to reflect the distance that non-bonded interactions have been shown to have measurable effects on the chemical shifts of protein^57,64^. We calculated the K_D_ value according to K_D_=P_u_/P_b_(*v*N_A_)^-1^ where (P_u_) is the fraction of frames with no ligand contacts, *v* is the volume of simulation box, *N*_Av_ is Avogadro’s number and *v*N_A_ is the simulated concentration^65^. Using these definitions, we observe that EPI-002 has a bound population of 43 ± 2% corresponding to a K_D_ value of 5.24 ± 0.43mM and EPI-7170 has a bound populations of 67 ± 2%, corresponding to a simulated K_D_ value of 1.92 ± 0.15mM. We report the simulated K_D_ values and estimated statistical errors as a function of simulation time in Fig. S19, illustrating that these quantities are well converged and that the difference in simulated K_D_ values is statistically significant.

We note that the absolute values of intermolecular contact probabilities and the corresponding K_D_ values will be sensitive to the distance thresholds used to define intermolecular contacts. These values are therefore most meaningfully compared to simulated K_D_ values calculated with the same distance thresholds and are not directly comparable to experimental K_D_ values measured from biophysical experiments. We and others^32^ have found that the distance threshold used to define intermolecular contacts has no effect on the ratios of K_D_ values and contact probabilities calculated between different ligands in multiple simulations. We note that in a previous study of small molecules binding to a 20-residue fragment of α-synuclein^32^ that used a more lenient 6.0Å contact threshold between all protein atoms (including hydrogens) and ligand heavy atoms, the lowest simulated K_D_ value observed among a series of 50 ligands was 4.5mM ± 0.15mM. The simulated affinity of EPI-002 to Tau-5_R2_R3_ is therefore on-par with the tightest binding ligands from that study, and the simulated K_D_ of EPI-7170 to Tau-5_R2_R3_ is ∼2.5-fold smaller than the tightest binding ligand from that study^32^.

### EPI-002 and EPI-7170 predominantly interact with aromatic residues at an interface between the R2 and R3 regions of Tau-5_R2_R3_

We observed that bound states of EPI-002 and EPI-7170 are highly dynamic and consist of a heterogenous ensemble of interconverting binding modes, consistent with the previously proposed *dynamic shuttling* mechanism^32^ (Movies S2-S3). This observation is consistent with the small magnitude of NMR CSPs observed in EPI-002 ligand titrations^24^. The contact probability of EPI-002 and EPI-7170 with each residue of Tau-5_R2_R3_ are shown in Fig. 2A. Convergence analyses of the per-residue contact probabilities are shown in Fig. S20-S21. We observe that EPI-7170 has an elevated contact probability with all residues relative to EPI-002 and that the most populated intermolecular contacts of both ligands are made with aromatic residues in the R2 and R3 regions.

**Figure 2.**
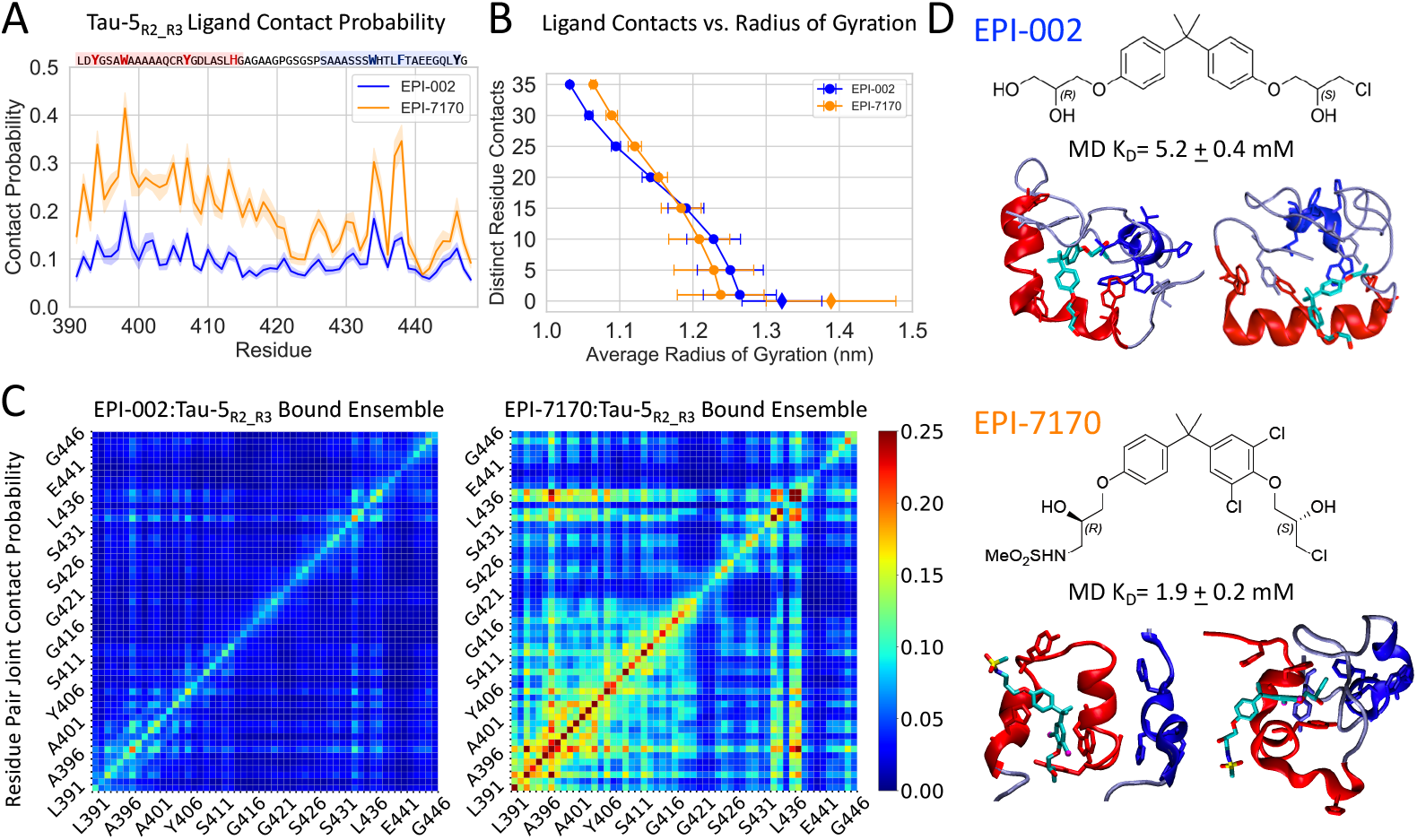
A) Per-residue contact probabilities observed between Tau-5_R2_R3_ and EPI-002 (blue) and EPI-7170 (orange) in REST2 MD simulations. Contacts are defined as occurring in frames where any non-hydrogen ligand atom is within 6.0Å of a non-hydrogen protein atom. Shaded regions indicate statistical error estimates from blocking. B) Average Cα-atom radius of gyratyion (R_g_) of Tau-5_R2_R3_ conformations as a function of the minimum number of distinct residue contacts formed in MD simulations. Error bars reflect the variance of R_g_ values for each subset of Tau-5_R2_R3_ conformations. Diamonds reflect the average radius of gyration of Tau-5_R2_R3_ conformations in frames with no protein ligand contacts in simulations in the presence of ligands. C) The probability that a pair of residues in Tau-5_R2_R3_ simultaneously form ligand contacts in the EPI-002:Tau-5_R2_R3_ and EPI-710:Tau-5_R2_R3_ bound ensembles. D) Chemical structures of EPI-002 and EPI-7170 and illustrative MD Snapshots of Tau-5_R2_R3_ interacting with ligands. The R2 and R3 helices of Tau-5_R2_R3_ are colored red and blue respectively. K_D_ values observed in MD simulations were calculated by defining the bound population (P_b_) as the fraction frames with at least one ligand contact.

The most populated contacts between EPI-002 and Tau-5_R2_R3_ are made by residues W397 and W433 and the most populated contacts between EPI-7170 and Tau-5_R2_R3_ are made by residues W397 and F437. We observed highly populated intramolecular contacts between these residue pairs in simulations of apo Tau-5_R2_R3_ (Fig. S1, Fig. S22). To determine if the EPI compounds are binding at an interface formed by the R2 and R3 regions we examined the probability of EPI-002 and EPI-7170 forming simultaneous contacts with each Tau-5_R2_R3_ residue pair (Fig. 2C). We observe substantially more cooperative binding to residues in both the R2 and R3 regions in the EPI-7170 bound ensemble. The bound ensembles of EPI-002 and EPI-7170 contain contacts with aromatic residues in both the R2 and R3 regions in 46.7% and 60.0% of bound conformations respectively. Our simulations therefore indicate that both EPI-002 and EPI-7170 predominantly bind at an interface between the R2 and R3 region of Tau-5_R2_R3_, and that EPI-7170 does so to a greater extent.

### EPI-002 and EPI-7170 binding induces the formation of collapsed helical molten-globule-like states of Tau-5_R2_R3_

In both EPI-002 and EPI-7170 bound ensembles, we found that bound conformations that form simultaneous contacts with greater numbers of Tau-5_R2_R3_ residues become progressively more compact (Fig. 2B) suggesting a scenario where the R2 and R3 regions can be thought to collapse around the ligands as they penetrate into the hydrophobic core of Tau-5_R2_R3_. We provide illustrative conformation of EPI-002 and EPI-7170 bound to more compact conformations Tau-5_R2_R3_ in Figure 2D, and Movies S2 and S3. These states, which do not have well defined hydrophobic cores and contain a heterogenous distribution of the locations and relative orientations of helical elements are consistent with the properties of partially folded molten-globule-like states observed protein folding^66-69^. Specifically, the ligand bound conformations of Tau-5_R2_R3_ are more compact than a random coil, and contain large clusters of aromatic and hydrophobic residues, but these hydrophobic cores are not rigid and well defined. The hydrophobic cores observed in different bound conformations Tau-5_R2_R3_ consist of many different combinations of residues and vary in size. Similarly, while there are substantially elevated populations of helical conformations in bound Tau-5_R2_R3_ conformations relative to random coil structures, the locations of these helical elements and their relative orientations differ substantially between conformations observed in the ligand bound ensembles.

In simulations of Tau-5_R2_R3_ in the presence of ligands, we observe that bound conformations of EPI-002 and EPI-7170 have average R_g_ values of 12.6 ± 0.3Å and 12.4 ± 0.2Å respectively, while the unbound conformations have average R_g_ values of 13.9 ± 0.4Å and 13.2 ± 0.2Å respectively (Fig 2B, Fig. S23). We note the unbound states of Tau-5_R2_R3_ from simulations in the presence of ligands have larger average R_g_ values than states observed in the apo simulation of Tau-5_R2_R3_ (12.5 ± 0.1Å). A direct comparison between these values is complicated by the higher affinity of EPI-002 and EPI-7170 to compact conformations of Tau-5_R2_R3_ (Fig. S23) and the effect that the simulated timescales of association and dissociation events of EPI-002 and EPI-7170 to Tau-5_R2_R3_ will have on the populations of extended states in the lower solute temperature replicas of REST2 simulations. As a large fraction of frames in the simulations of Tau-5_R2_R3_ in the presence of ligands are bound, the apo simulation of Tau-5_R2_R3_ more extensively samples unbound states that are unperturbed by transient interactions with ligands. We therefore expect that the conformational properties of Tau-5_R2_R3_ observed in the apo REST2 MD simulation provide a more relevant comparison to the conformational properties of the ligand bound ensembles.

To better understand the effects of ligand binding on the conformations of Tau-5_R2_R3_ we compare the helical propensity, R_g_ and the Sα helical order parameters of the EPI-002 and EPI-7170 bound states to the apo Tau-5_R2_R3_ trajectory (Table 1, Fig. 3). We observe that differences in the per-residue helical fractions of the apo Tau-5_R2_R3_ ensemble and the EPI-002:Tau-5_R2_R3_ bound ensemble are largely within statistical error estimates but the per-residue helical fractions of the EPI-7170:Tau-5_R2_R3_ bound ensemble are substantially higher than the apo and EPI-002 bound ensembles. We observe more pronounced differences between the Tau-5_R2_R3_ apo ensemble and the ligand bound ensembles when considering differences in the Sα order parameter, which report on the cooperative formation of longer helical elements and the simultaneous formation of helical elements distant in sequence (See Methods). The free energy surface of the apo and ligand bound ensembles of Tau-5_R2_R3_ are shown as a function of Sα (Fig. 3B) and as a function of R_g_ and Sα (Fig. 3C). The apo ensemble of Tau-5_R2_R3_ has a pronounced free energy minimum centered at (Sα=5, R_g_=1.2nm), a shallower minimum centered at (Sα=9, R_g_=1.1nm) and no substantial minima with Sα>10. The EPI-002 bound ensemble has a broad free energy minima centered at (Sα= 7.5, R_g_=1.1nm) and in contrast to the apo ensemble, has an additional minimum ∼1kcal/mol higher in free energy centered at an (Sα= 20.0, R_g_=1.1nm). The EPI-7170 ensemble is globally shifted to higher Sα regions relative to both ensembles, with a global minimum centered at (Sα= 9.0, R_g_=1.1nm) and a substantially elevated population of conformations with Sα >10.0. This indicates that EPI-002 and EPI-7170 both stabilize the cooperative formation helical conformations in the R2 and R3 regions of Tau-5_R2_R3_, though EPI-7170 does so to a greater extent.

**Figure 3.**
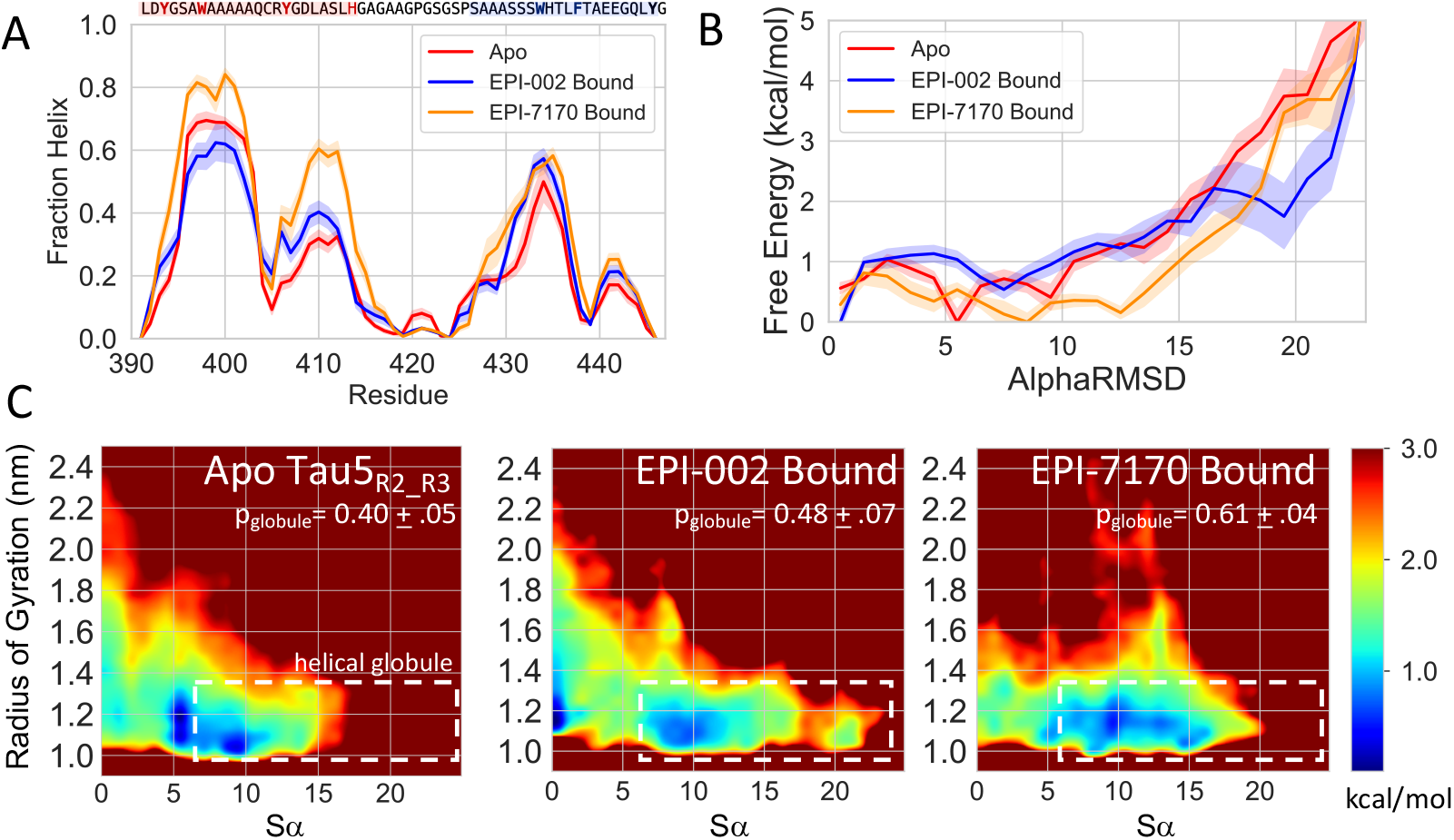
A) Helical Propensities observed in the 300K replica of explicit solvent REST2 MD simulations of Tau-5_R2_R3_ in its apo form (red) and in bound conformations obtained from simulations run in presence of EPI-002 (blue) and EPI-7170 (orange). B) Free energy surface of Tau-5_R2_R3_ conformations at 300K as a function of the helical collective variable Sα for each ensemble. Sα describes the similarity of all consecutive 6-residue fragments to ideal helical geometries. 6-residue fragments with an RMSD<0.8Å from a canonical helix contribute a value of 1 to Sα while 6-residue fragments with an RMSD>4.0 Å contribute a value of 0 to Sα. Shaded regions indicate statistical error estimates from blocking. C) Free energy surfaces as a function of radius of gyration (R_g_; reported in nm) and Sα. The dotted white lines indicate the defined boundary of “helical globule” state (Sα> 6.0, R_g_<1.3nm). The population of the helical globule state is reported as p_globule_.

In order to quantify the relative populations of collapsed helical conformations in each ensemble we define a “helical globule” state in each ensemble as all Tau-5_R2_R3_ conformation with Sα >6.0 and Rg<1.3nm and report the populations and relative free energies (ΔG_globule_) at 300K of this state in each simulation (Table 2, Figure 3C). Conformations with Sα >6.0 correspond to Tau-5_R2_R3_ conformations where at least 6 6-residue fragments have small RMSDs from an ideal helical conformation (See Methods). We observe that the helical globule state has a population of 40.4 ± 5.2% in the apo simulation, a modestly increased population in the EPI-002 bound ensemble (48.5 ± 6.5%) and a substantially larger population in EPI-7170 bound ensemble (61.1 ± 4.5%). We note that relative populations of this state are sensitive to the selected threshold of Sα. We selected the threshold value of Sα=6.0 based on our desire to quantify the stability of the ligand bound states with more cooperative helix formation than is observed in the global free energy minima of the apo ensemble. We compare the calculated populations of the helical globule state as a function of the selected cutoff value of Sα in Fig. S24. We observe that the helical globule populations of ligand bound states increase relative to the helical globule populations of the Tau-5_R2_R3_ apo ensemble as larger values Sα are used as a cutoff.

### Aromatic stacking interactions of the dichlorinated phenyl ring of EPI-7170 stabilize compact helical conformations of Tau-5_R2_R3_

We conducted a detailed dissection of the intermolecular interactions between Tau-5_R2_R3_ and the EPI compounds to better understand the molecular features of EPI-7170 that confer tighter binding and stabilize collapsed helical conformations of Tau-5_R2_R3_. The populations of intermolecular hydrophobic contacts, hydrogen bonds, and aromatic stacking interactions with each residue of Tau-5_R2_R3_ are shown for both ligands in Fig. 4 (See Methods). We observe relatively similar hydrophobic contact profiles between the two ligands, with the exception of elevated hydrophobic contact probabilities between EPI-7170 and ^436^LF^437^.

**Figure 4.**
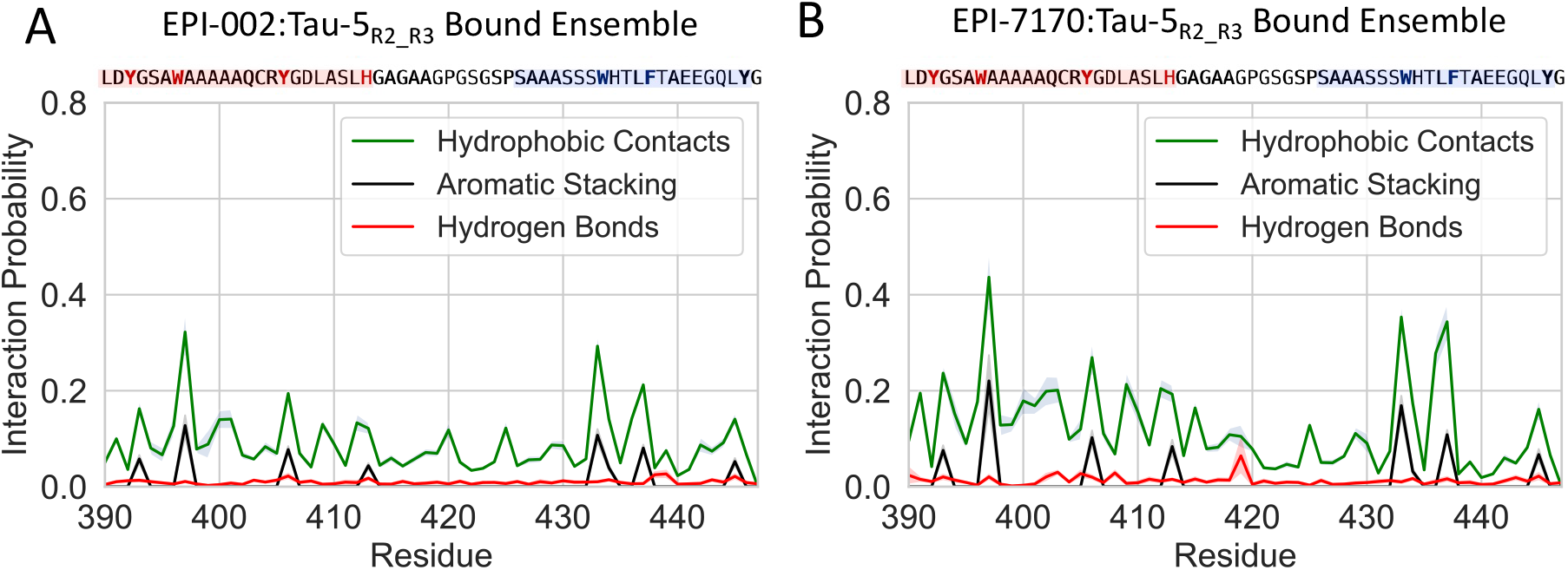
Populations of intermolecular interactions observed in the Tau-5_R2_R3_:EPI-002 and Tau-5_R2_R3_:EPI-7170 bound ensembles. Populations are calculated considering only the bound frames of MD simulations in the presence of ligands. Shaded regions indicate statistical error estimates from blocking.

The most significant differences in the binding modes of EPI-002 and EPI-7170 are the elevated populations of aromatic stacking interactions in the EPI-7170 bound ensemble (See Methods). The stacking populations of each aromatic residue in the EPI-002 and EPI-7170 bound ensembles are shown in Fig. 5A. The largest increases in stacking propensity in the EPI-7170 bound ensemble relative to the EPI-002 bound ensemble occur in residues W397 (15% vs 5% population) and W433 (11% vs 5%). All aromatic residues experience a 2-3-fold increase in stacking propensity in the EPI-7170 bound ensemble relative to the EPI-002 bound ensemble. We examined the orientations of the stacking interactions occurring in both ensembles, calculating the propensity of each ligand ring to form face-to-face parallel stacking interactions and T-shaped stacking interactions with each aromatic residue (Fig. 5, Fig. S25-S26). We find that the dichlorinated phenyl ring of EPI-7170 forms substantially more parallel stacking interactions than the unchlorinated EPI-7170 phenyl ring and both phenyl rings of EPI-002 (Fig. S25-26). We also observe that the unchlorinated phenyl rings of EPI-002 and EPI-7170 sample a broader distribution of ring orientations than the chlorinated ring of EPI-7170, with fewer well-defined free energy minima.

**Figure 5.**
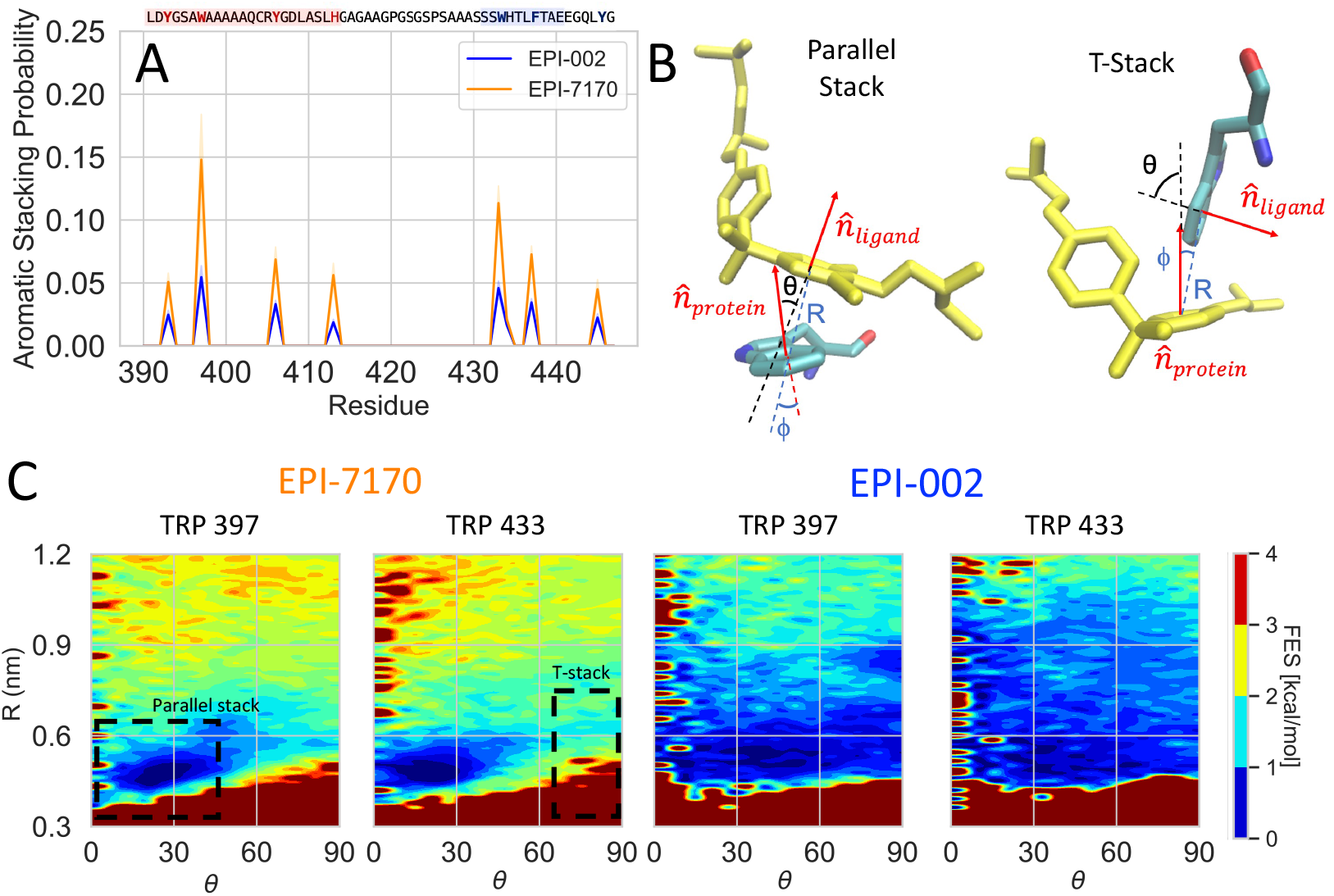
A) Populations of aromatic stacking contacts between aromatic sidechains of Tau-5_R2_R3_ and aromatic rings of EPI-002 and EPI-7170 observed in the 300K replica of explicit solvent REST2 MD simulations. Shaded regions indicate statistical error estimates from blocking. B) Definition of angles and distances used to define stacking orientations and examples of parallel stacked and T-stacked conformations between EPI compounds and a tyrosine sidechain. For a protein aromatic ring and a ligand aromatic ring R is defined as the distance between ring centers, 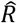 is defined as unit vector connecting the ring centers, 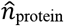 and 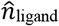 are normal vectors to the sidechain and ligand ring planes originating from the ring centers, θ is the angle between 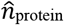 and 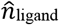, and ϕ is the angle between 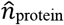 and 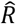. Parallel stacked conformations are defined as occurring when R<6.5Å, θ < 45° & ϕ < 60° and T-stacked conformations are defined as occurring when R<7.5Å, θ > 75° & ϕ < 60°. C) Free energy surfaces as a function of R and θ between the chlorinated phenyl ring of EPI-7170 and the equivalent non-chlorinated phenyl ring in EPI-002 and the indole rings of TYR 397 and TYR 433.

We observe an ∼2-fold increase in hydrogen bond population between EPI-7170 and the R2 residues ^403^QCRYGD^408^ relative to the hydrogen bond populations observed in the EPI-002 bound ensemble. The average hydrogen bond propensity for these residues is 2.1 ± 0.4% in the EPI-7170 bound ensemble compared to 1.3 ± 0.2% in the EPI-002 bound ensemble (Fig 4, Fig S27). The most populated hydrogen bond interactions occur with Q403, R405 and D408. The populations of Q403, R405 and D408 hydrogen bonds are 3.0 ± 0.3%, 2.7 ± 0.6% and 2.9 ± 0.5% in the EPI-7170 bound ensemble and 1.4 ± 0.2%, 1.4 ± 0.2% and 1.2 ± 0.2% in the EPI-002 bound ensemble, respectively. For each of these residues there are many distinct hydrogen bond pairs with similar populations. These hydrogen bonds contain both sidechain and backbone atoms functioning as donors and acceptors with different ligand atoms in the alkyl chains of EPI-002 and EPI-7170. In general, we cannot identify small subsets of dominant hydrogen bonding pairs in any region of Tau-5_R2_R3_ with the exception of a relatively highly populated hydrogen bond between the backbone amide of G419 and the oxygen atom in the chlorohydrin group of EPI-7170 (6.4 ± 3.4%). This hydrogen bond is predominantly populated in a metastable set of bound conformations sampled in a contiguous 900ns portion of the trajectory, and correspondingly its population has a larger statistical error estimate than all other hydrogen bond populations. W397 hydrophobic contacts and aromatic stacking interactions are formed in the majority of bound frames containing this hydrogen bond (95% and 75% respectively), illustrating that this interaction is predominantly present in a relatively narrow subset of bound conformations compared to more dynamic intermolecular interactions that are observed in more conformationally diverse subensembles of bound conformations.

We observe that residues with the largest hydrogen bond propensities (Q403, R405, D408) all have relatively high conditional interaction probabilities with hydrophobic and aromatic stacking interactions in other regions of Tau-5. W397 hydrophobic interactions occur in ∼50% of frames containing hydrogen bonds between EPI-7170 and Q403, and W397 aromatic stacking interactions occur in ∼25% of frames containing hydrogens bonds with Q403. F437 hydrophobic and aromatic stacking interactions occur in ∼50% and ∼25% of frames containing hydrogen bonds between EPI-7170 and R405, respectively, and Y445 hydrophobic and aromatic interactions occur in ∼70% and ∼50% of frames containing hydrogen bonds between EPI-7170 and D408, respectively. These high conditional interaction probabilities suggest the increased hydrogen bond populations observed in the EPI-7170 bound ensemble relative to the EPI-002 bound ensemble result from the dichlorinated phenyl ring of EPI-7170 more effectively localizing the ligand in the aromatic core of Tau-5_R2_R3_ through its increased hydrophobicity and planar aromatic stacking propensity. Once EPI-7170 is buried in helical globule states of Tau-5_R2_R3_ more hydrogen bonds likely become accessible through small displacements in ligand positions via a dynamic shuttling mechanism^32^. We note that the conditional interaction probabilities between hydrogen bonds and hydrophobic and aromatic interactions with non-neighboring residues are substantially higher than those observed in simulations of lower affinities compounds of α-synuclein^32^, suggesting that the more cooperative formation of intermolecular interactions observed in the EPI-002 and EPI-7170 bound ensembles confer the higher affinity binding observed in this investigation.

## Discussion

The design of small molecule inhibitors targeting the disordered N-terminal transactivation domain (NTD) of the androgen receptor is a promising avenue for the development of drugs to treat castration-resistant prostate cancer (CPRC) and an important proof-of-principle of the feasibility of IDP drug design. Recent preclinical results for EPI-7386^46^, a second-generation AR-NTD inhibitor from the EPI-7170 compound family which entered human trials in 2020^45^, are encouraging and suggest that EPI family compounds are viable CRPC drug candidates. A large amount of biophysical and biological research has been carried out to characterize the interactions of EPI compounds with the AR-NTD and their therapeutic potential, but until now, no atomic resolution structural models have existed to explain the molecular mechanism of AR-NTD inhibition or rationalize differences in the activity of compounds in the EPI family.

In this investigation, we have leveraged recent advances in molecular simulation force fields and enhanced sampling methods to provide atomic resolution binding mechanism models of EPI-002 and EPI-7170 that are highly consistent with previously reported NMR measurements^24^. We utilized long-timescale enhanced sampling MD simulations with the state-of-the-art a99SB-disp force field^49^ to study the conformational properties of Tau-5_R2_R3_, a disordered fragment of the AR-NTD that contains the Tau-5 residues that showed the most pronounced NMR chemical shift perturbations in EPI-002 titrations^24^. We found that the apo ensemble of Tau-5_R2_R3_ obtained from MD simulations is in excellent agreement with previously measured NMR chemical shift data^24^. Quantitative comparisons with backbone chemical shifts reveal relatively minor discrepancies in the simulated helical propensity of the R3 region, which can be corrected using a maximum-entropy ensemble reweighting algorithm^61,62^. The apo ensemble of Tau-5_R2_R3_ reveals substantially populated intramolecular contacts between the R2 and R3 regions, which were previously hypothesized to be important for EPI-002 binding^24^. Simulations of Tau-5_R2_R3_ in the presence of EPI-002 revealed that EPI-002 binds via a dynamic and heterogenous ensemble of interconverting binding modes and does not induce the formation of a stable folded conformation of Tau-5_R2_R3_, an observation that is consistent with relatively small NMR CSPs observed in ligand titrations^24^. We find that EPI-002 binding is predominantly driven by interactions with aromatic residues in the transiently formed R2 and R3 helices, and that these helical regions can “wrap around” EPI-002 to form compact helical conformations that resemble dynamic molten-globe states from protein folding, which we define as a helical globule state.

The compound EPI-7170 has a 4-6-dichloro substituted phenyl ring in its bisphenol-A scaffold and a methylsulfonamide group in place of a hydroxyl group in an alkyl chain relative to EPI-002. MD simulations of Tau-5_R2_R3_ in the presence of EPI-7170 reveal that EPI-7170 has an ∼2.5-fold higher affinity to Tau-5_R2_R3_ than EPI-002. We observe that EPI-7170 has a substantially higher probability of simultaneously interacting with residues in both the R2 and R3 helices in its bound ensemble. EPI-7170 binding substantially increases the helical populations of Tau-5_R2_R3_ relative to the apo Tau-5_R2_R3_ ensemble and the EPI-002:Tau-5_R2_R3_ bound ensemble. Analysis of the α-helical order parameter (Sα) reveals that EPI-002 and EPI-7170 both induce the cooperative formation of helical elements in the R2 and R3 regions and that the EPI-7170:Tau-5_R2_R3_ bound ensemble has a substantially elevated population of helical globule conformations relative to the apo Tau-5_R2_R3_ ensemble and EPI-002:Tau-5_R2_R3_ bound ensemble. Our simulations identify that the increased affinity of EPI-7170 results from an increased propensity of the dechlorinated phenyl ring to form parallel face-to-face stacking interactions with aromatic sidechains relative to the phenyl rings EPI-002. These stacking interactions localize EPI-7170 to a dynamic hydrophobic/aromatic core of Tau-5_R2_R3_, where it adopts an increased propensity to form an array of interconverting hydrogen bonding interactions with residues in the R2 region.

NMR measurements of the interaction between EPI-7170 and Tau-5 have yet-to-be reported, which may be the result of the relative insolubility of EPI-7170 relative to EPI-002^42^. A number of small molecule ligands that bind IDPs have been found to be relatively insoluble at concentrations required for biophysical experiments, in particular NMR^32^. This suggests that solubility may be a ubiquitous problem when studying small molecules that bind IDPs. This underscores the value of combining insights from experimental and computational studies when studying compounds near the detectable solubility limits of biophysical experiments. By validating our simulations of apo Tau-5_R2_R3_ and EPI-002 Tau-5_R2_R3_ binding against NMR data, we gain confidence in the ability of our simulation protocol to describe the conformational properties of Tau-5_R2_R3_ and the interactions between Tau-5_R2_R3_ and EPI compounds with a common bisphenol-A scaffold. Given the relatively small differences in the chemical structures of EPI-002 and EPI-7170 and quality of ligand molecular mechanics force fields^32,63^, we expect that our simulation model will be capable of discerning differences in their binding modes as observed in a previous study^32^.

While NMR data has not been reported for interactions between EPI-7170 and Tau-5_R2_R3_, protein and ligand-detected NMR data has been reported for the related compound EPI-7386^46^. These studies have identified NMR CSPs of the sidechain resonances of W397 and W433 of Tau-5 in EPI-7386 titrations, which is consistent with the importance of these residues for EPI-7170 binding in our simulations. Saturation Transfer Difference (STD) experiments also confirm direct interaction between EPI-7386 and Tau-5, demonstrating that these CSPs do not only result from allosteric changes in the conformational ensemble of Tau-5 upon EPI-7386 binding^46^.

Molecular simulation studies of the interactions between small molecules and IDPs are becoming more common^30,32-36^, but simulation studies that provide detailed comparisons of the binding modes of multiple small molecules and identify subsets of intermolecular interactions that confer differences in their specificity and affinity have only recently begun to emerge^32^. Understanding the molecular features of dynamic ligand binding modes that confer affinity and specificity among known IDP ligands is an essential step in developing rational design strategies to improve the affinity of small molecules to IDPs or to design inhibitors *de novo* for new IDP sequences. Identifying the key intermolecular interactions that underpin IDP ligand binding modes enables medicinal chemists to identify promising regions of chemical space to explore for further ligand derivatization and provides experimentally testable hypotheses to validate and refine binding mechanism models in drug discovery campaigns. In this investigation, we have identified the importance of aromatic stacking orientations of the phenyl groups of the Bisphenol-A scaffold and hydrogen bonding interactions in the alkyl chains of EPI compounds for stabilizing compact helical globule states of Tau-5_R2_R3_. These mechanistic insights suggest important small molecule features to explore in the development of more potent androgen receptor inhibitors for the treatment of CRPC.

The simulations conducted in this study were carried on a fragment of the Tau-5 region of AR containing the residues previously observed to have the highest propensity to interact with EPI-002 to obtain a computationally tractable system and enable statistically meaningful comparisons of the binding modes of EPI-002 and EPI-7170. We note that the R1 helix of Tau-5 was also found to interact with EPI-002 by NMR in the context of the full Tau-5 region but that a peptide containing only the R1 domain was not found to interact with EPI-002 at similar concentrations. Based on the sequence composition of R1, in particular the presence of 6 aromatic residues in a 31 residue domain and the helical propensity observed by NMR, we speculate that EPI-002 and EPI-7170 likely interact with the with R1 region with similar bindings mechanisms to those observed in this investigation. We hypothesize that EPI compounds stabilize transient long-range intramolecular between the R1 and R2 regions, R1 and R3 regions, and simultaneous interactions between the R1, R2 and R3 regions with similar binding modes to those observed between the R2 and R3 regions in this investigation. Due to the presence of the relatively rigid polyproline linker between R1 and R2, we suspect that interactions involving the R1 helix are less important for conferring the affinity of EPI compounds to Tau-5 than the interactions between R2 and R3 described here. Studying the interactions of EPI compounds with larger constructs of Tau-5 that contain the R1, R2 and R3 regions will require substantially longer simulations than those reported here, and in practice will likely require the optimization of additional enhanced sampling algorithms for this system.

Finally, we note that the chlorohydrin group of EPI-002 has been found to be weakly covalently reactive with Tau-5, and it is hypothesized that covalent attachment of EPI-002 to Tau-5 may be important for its biological activity^14,37^. The atomically detailed bound ensembles of EPI-002:Tau-5_R2_R3_ and EPI-7170:Tau-5_R2_R3_ characterized in this investigation will enable atomic resolution comparisons between non-covalent and covalent binding and inhibition mechanisms of EPI compounds in future simulation studies.

## Methods

### Simulation Methods

Simulations of androgen receptor Tau-5_R2_R3_ region (residues L391-G446, capped with ACE and NH2 groups) in the presence and absence of ligands were performed using GROMACS 2019.2^70-72^ patched with PLUMED v2.6.0^73^. The AR Tau-5_R2− R3_ protein was parameterized using the a99SB-*disp* force field and water molecules were parameterized with the a99SB-disp water model^49^. Simulations with ligands were run using GAFF1^63^ for ligand forcefield parameters, obtained from ACPYPE^74^. Each system was solvated with 13200 water molecules in a cubic box with a length of 7.5nm and neutralized with a salt concentration of 20 mM NaCl by 8 Na^+^ ions and 5 Cl^−^ ions. Energy minimization of the system is performed with steepest descent minimization algorithm to the maximum force smaller than 1000.0 kJ/(mol/nm). Equilibration was first performed in NVT ensemble for 2000 ps at the temperature of 300 K using the Berendsen thermostat^75^; then the systems were further equilibrated in NPT ensemble for 200 ps at a target pressure of 1 bar with the temperature at 300 K maintained by Berendsen thermostat, with position restraints added to all heavy atoms. Bond lengths and angles of protein and ligand atoms were constrained with the LINCS^76^ algorithm and water constraints were applied using the SETTLE algorithm^77^. Canonical sampling in the NVT ensemble algorithms was obtained using the Bussi et. al. velocity rescaling thermostat^78^ with a 2 femtosecond timestep. The PME algorithm^79^ was utilized for electrostatics with a grid spacing 1.6nm. Van der Waals forces were calculated using a 0.9nm cut-off length. The REST2 algorithm^53,80^ was utilized with exchange attempted every 80ps, selecting all protein atoms as the solute region. A 16-replica temperature ladder ranging from 300-500K was utilized to scale the solute temperature. Apo Tau-5_R2− R3_, Tau-5_R2− R3_ + EPI-002, Tau-5_R2− R3_ + EPI-7170 were simulated for 4.6μs, 4.0μs and 4.5μs per replica respectively, for total simulation times of 74μs, 64μs and 72μs respectively. Frames were saved every 80ps for analysis.

Initial structures of Tau-5_R2_R3_ were generated with the pmx software^81^. Pmx builder was used to generate a fully helical Tau-5_R2_R3_ conformation with all residues in an ideal helical conformation (ϕ=-57,Ψ=-47) and a partially helical Tau-5_R2_R3_ conformation where only residues in the R2 and R3 regions were built in helical conformations and the remaining residues were left in an extended conformation. These starting structures were subject to an energy minimization, and then each structure was used to perform a short 100ps 600K high temperature unfolding simulation in vacuo in the NVT ensemble. 8 structures were selected from each of the unfolding trajectories to span a range of helicity in starting structures. Initial conformations of EPI-002 and EPI-7170 were generated using the Open Babel online toolbox^82^. Starting structures for Tau-5_R2_R3_ simulations in the presence of ligands were prepared by inserting the ligands into simulation boxes with the same 16 Tau-5_R2_R3_ starting structures and repeating the process of solvation, neutralization, energy minimization and NVT and NPT equilibration as described above.

### Statistical Error Estimates

Statistical error estimates of the simulated properties from MD simulations were calculated using a blocking analysis following Flyvbjerg and Peterson^55^ with an optimal block size selection determined as described by Wolff^56^, using the *pyblock* python package. In this procedure, the trajectory is divided into a given number of equally sized “blocks”, average values of simulated quantities are computed for each block, and the standard error of the average values calculated across all blocks is used as an error estimate. An optimal block size is selected to minimize the estimated error of the standard error across blocks according to Wolff^56^.

### Maximum-Entropy Ensemble Reweighting

NMR chemical shifts were calculated for the 300K base replica of the REST2 MD simulation of apo Tau-5_R2_R3_ using SPARTA+^57^. We utilized the maximum entropy reweighting algorithm of Cesari et. al^61,62^ with Cα NMR chemical shifts as restraints. We utilized a gaussian error model for Cα chemical shift predictions with a standard deviation σ=1.73ppm. The Kish ratio^83,84^ of the resulting ensemble was 43.9, which means that the algorithm effectively retained 43.9% of the frames from the unbiased simulation in the reweighted ensemble.

### Sα α-helical order parameter

The α-helical order parameter *S*α, measures the similarity of all 6-residue segments to an ideal helical structure (ϕ=-57,Ψ=-47)^54^. *S*α is calculated according to:

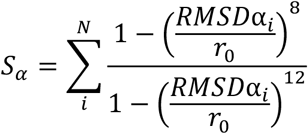

where the sum is over *N* consecutive 6-residue segments, RMSDα_*i*_ is the Cα-RMSD between an ideal α-helical geometry a 6-residue fragment (spanning from residue *i* to residue *i*+5), and *r*_0_ =0.8 Å. When *r*_0_ = 0.8 Å, a 6-residue fragment with a value of RMSD_α_ < 0.4 Å contributes a value of ∼1 to the *S*_α_ sum, a 6-residue fragment with a value of RMSD_α_=0.9 contributes a value of ∼0.5 to the *S*_α_ sum, and a 6-residue fragment with a value of RMSD_α_ > 2.5 Å contributes a value of ∼0 to the *S*_α_ sum. The value of *S*_α_ for a protein conformation can therefore be interpreted as proxy for the number of 6-residue fragments closely resembling an ideal helical conformation. A completely helical conformation of the 56 residue Tau-5_R2_R3_ construct has an *S*_α_ value of 51, and a Tau-5_R2_R3_ with no helical content has an *S*_α_ value of 0.

### Ligand Contacts and K_D_ calculations

We define an intermolecular contact between a ligand and a protein residue as occurring in any frame where at least one heavy (non-hydrogen) atom of that residue is found within 6.0Å of a ligand heavy atom. To calculate a simulated K_D_ value for each compound, we define the bound population (P_b_) of each ligand as the fraction of frames with at least one intermolecular contact between a ligand and Tau-5_R2_R3_. We calculated the K_D_ value according to K_D_=P_u_/P_b_(*v*N_A_)^-1^ where (P_u_) is the fraction of frames with no ligand contacts, *v* is the volume of simulation box and *N*_Av_ is Avogadro’s number number^65^. In the 7.5nm simulation box used in this work containing 1 ligand and 1 protein molecule, *v=*4.22*10^−25^L and the concentration of the ligand and protein are each 3.93mM.

### Ligand Intermolecular Interactions

Intermolecular hydrophobic contacts were defined as occurring when pairs of protein carbon and ligand carbon and protein carbon and ligand chlorine atoms were within 4Å. Potential hydrogen bond donors were defined as all nitrogen, oxygen, or sulfur atoms with an attached hydrogen, and potential hydrogen bond acceptors were defined as all nitrogen, oxygen and sulfur atoms. Hydrogen bonds were identified with a distance cutoff of 3.5 Å between the donor hydrogen and heavy-atom acceptor, and a donor-hydrogen-acceptor angle >150°.

Aromatic stacking interactions were calculated following the conventions Marsili et al.^85^ (Shown in Fig. 5) with modified distance and angle cutoffs. The cutoffs were selected based on the observed locations of free energy minima of parallel (face-to-face) stacked and T-stacked conformations between the EPI ligands and protein aromatic sidechains. For a protein aromatic ring and a ligand aromatic ring: we define R as the distance between ring centers, 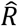 as the unit vector connecting the ring centers, 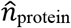 and 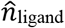 are normal vectors to the sidechain and ligand ring planes originating from the ring centers, θ is the angle between 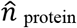 and 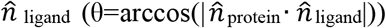, and ϕ is the angle between 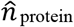 and 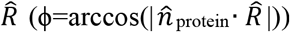. Parallel stacked conformations were defined as occurring when R<6.5Å, θ < 60° & ϕ < 45° and T-stacked conformations were defined as occurring when R<7.5Å, θ > 75° & ϕ < 45°. In the evaluation of the free energy surfaces, we chose not to distinguish between angles of θ and π– θ and between angles of ϕ and (π–ϕ). Distance–angle probability distributions and corresponding free energy surfaces (Fig. 5) were normalized by a factor *R*^2^ sin(θ) to obtain a flat free energy profile in the case of configurations with no angle or distance preference^85,86^.

## Supporting Information

Convergence Analysis; Figures S1-S27; Movies S1-S3.

## Data Availability

All trajectories reported in this investigation and code used for analyses are freely available from https://github.com/paulrobustelli/AR_ligand_binding.

## Acknowledgements

This work was supported by the National Institutes of Health under award R35GM142750, the National Research Council supercomputing grant MCB200087P, AGAUR (2017 SGR 324), MINECO (PID2019-110198RB-I00) and the European Research Council (CONCERT, contract number 648201). Jiaqi Zhu was supported by the China Scholarship Council and R35GM142750. The authors thank Massimiliano Bonomi for assistance implementing maximum-entropy reweighting methods, Matteo Paloni for providing scripts to calculate stacking interactions, and Stase Bielskute and Borja Mateos for stimulating discussions and a critical reading of this manuscript. IRB Barcelona is the recipient of a Severo Ochoa Award of Excellence from MINECO.

## Ethics Declarations

Paul Robustelli is a scientific advisor of Dewpoint Therapeutics and an Open Science Fellow and scientific consultant of Roivant Sciences. Xavier Salvatella is a founder and board member of Nuage Therapeutics. These affiliations have not influenced this work. The remaining authors declare no competing interests.

