## Supplemental Information for "Small Molecules Targeting the Disordered Transactivation Domain of the Androgen Receptor Induce the Formation of Collapsed Helical States"

Paul Robustelli

**MD Simulation Convergence Analyses**

Convergence of the Tau-$5_{R2\_R3}$ REST2 MD simulation was assessed by a comparison of the secondary structure profiles, free energy surfaces of Tau-$5_{R2\_R3}$ conformations as a function of the alpha helical order parameter S$\alpha$ (see main text methods) and the radius of gyration ($R_g$) and the intramolecular contact probabilities for each solute temperature rung in the REST2 temperature ladder (Fig. S1-S3). The relatively smooth solute-temperature dependence of each of these properties suggests the simulations are reasonably well converged. The same analyses were also performed on this simulation using demultiplexed replicas, which follow each independent replica through temperature space to determine if any individual replicas became stuck in local minima as they diffuse through the temperature ladder (Fig. S4-S6). We also found the statistical fluctuations to be relatively well converged among demultiplexed replicas; with only 3 of replicas (replicas 6, 11 and 12) showing substantial deviations from the average contact probabilities and secondary structure profiles relative to the remaining 13 replicas.

The helical propensity, intramolecular contact probabilities, and free energy surfaces of each replica as a function of $R_g$ and the $\alpha$-helical order parameter S$\alpha$ are shown for each temperature replica of a REST2 simulation of Tau-$5_{R2\_R3}$ in the presence of EPI-002 in Fig. S7-S9 and for each demultiplexed replica in Fig. S10-S12. We observed that the helical propensities decrease relatively smoothly and continuously across temperature replicas as the solute temperature increases, and the demultiplexed replicas have similar helical propensity profiles, with helical elements localized to the same region and differing only in magnitude between replicas. The helical propensity, contact probabilities, and free energy surfaces of each replica are shown for each temperature replica of a REST2 simulation of Tau-$5_{R2\_R3}$ in the presence of EPI-7170 in Fig. S13-S15 and for each demultiplexed replica in Fig. S16-S18. The smoothly varying $\alpha$-helical and $\beta$-sheet propensities across temperature replicas and the similarity of helical propensities, intramolecular contact probabilities, and free energy surfaces as function of $R_g$ and S$\alpha$ of the demultiplexed replicas suggest that simulations of Tau-$5_{R2\_R3}$ + EPI-7170 are exceptionally well converged.

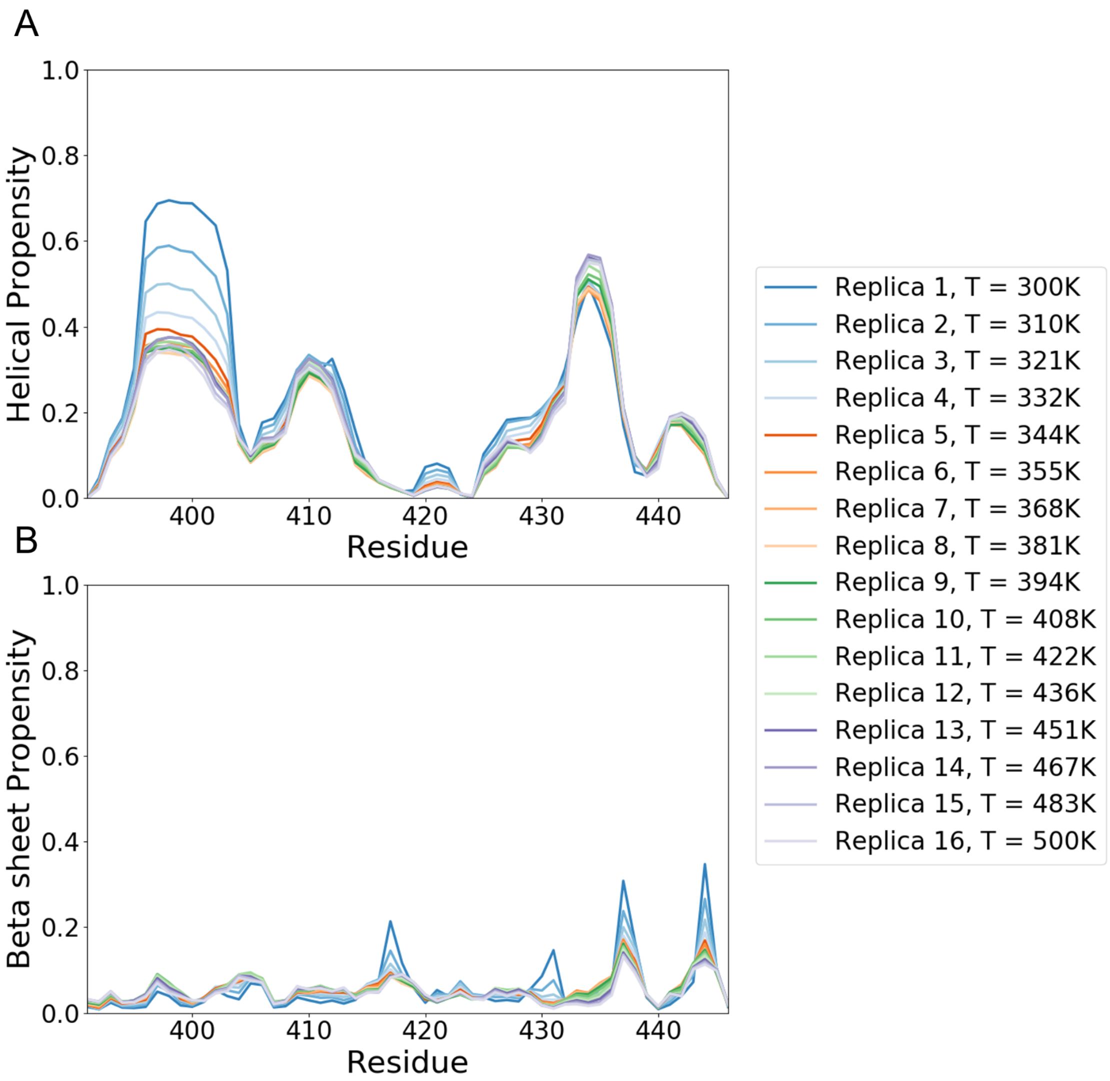


**Figure S1**. Comparison of α-helical (A) and β-sheet (B) propensities observed in the 16 temperature rungs of an apo Tau-$5_{R2\_R3}$ REST2 MD simulation. Secondary structure content is calculated by the DSSP algorithm. The solute temperature of each replica is shown in the figure legend.

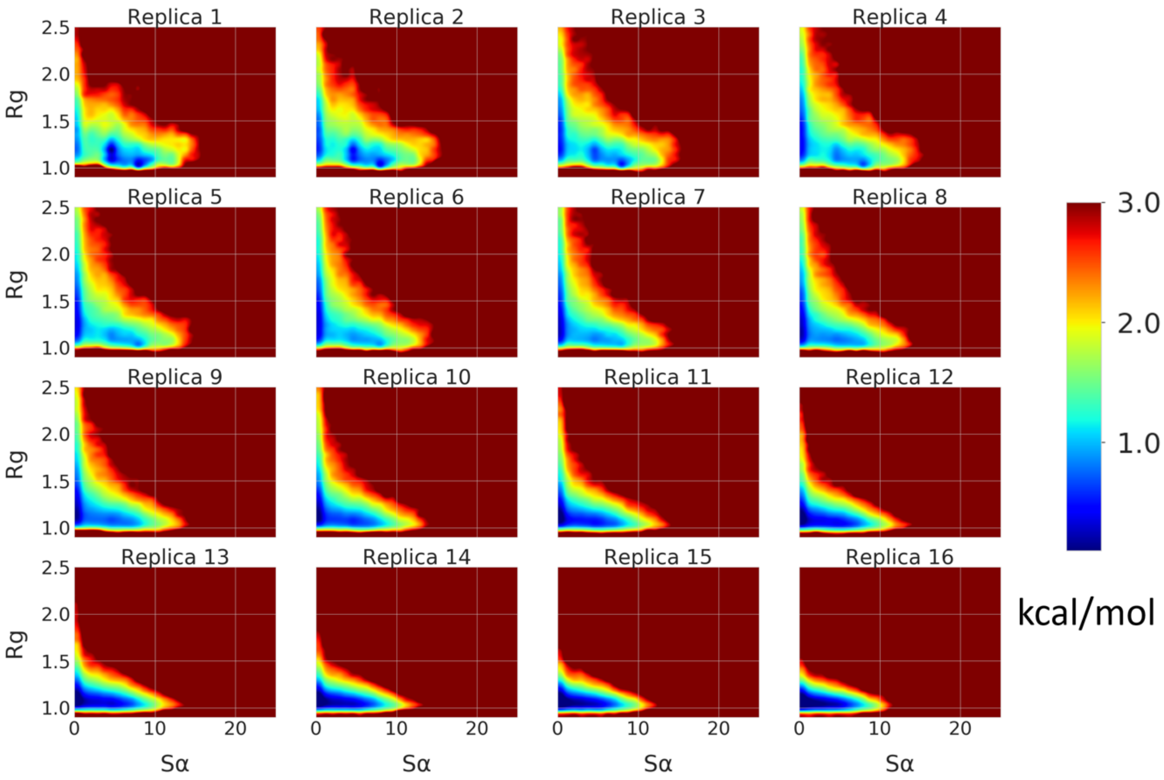


**Figure S2**. Comparison of free energy surfaces of the 16 solute temperature rungs of an apo Tau-$5_{R2\_R3}$ REST2 MD simulation as a function of the α-helical order parameter Sα and radius of gyration ($R_g$). $R_g$ is reported in nm.

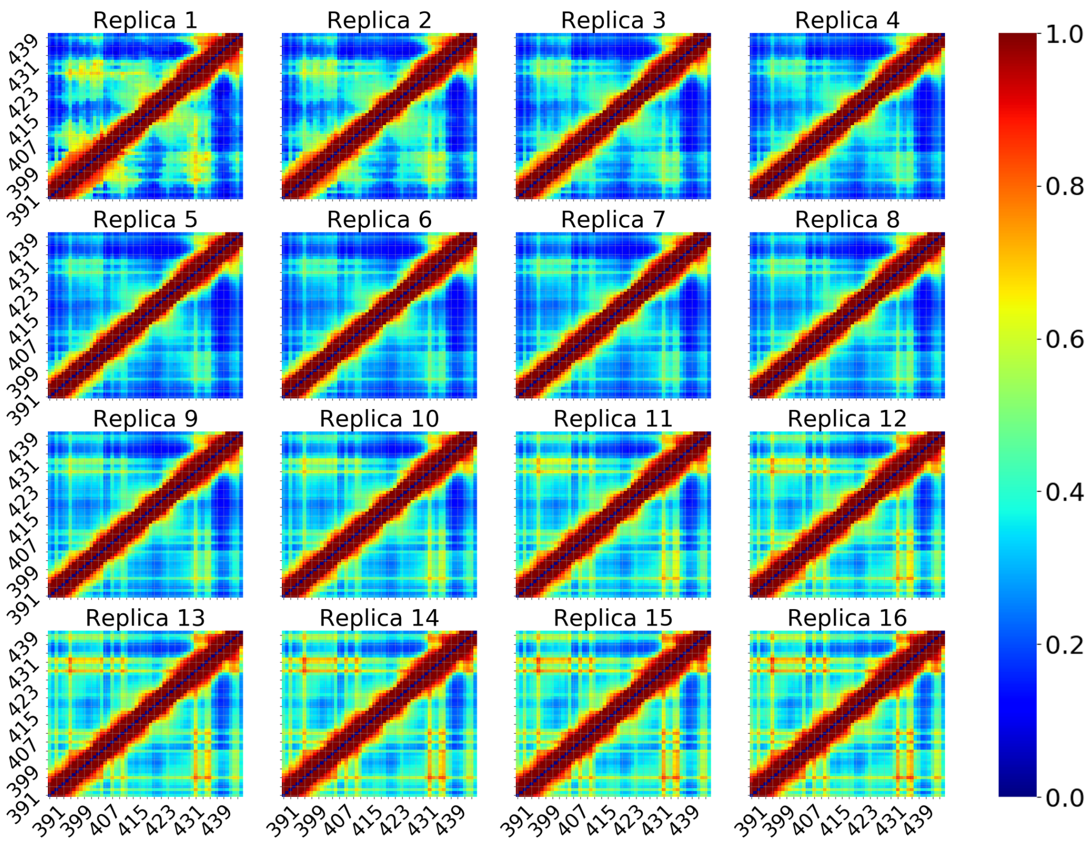


**Figure S3**. Comparison of the intramolecular contact probabilities observed in the 16 solute temperature rungs of an apo Tau-$5_{R2\_R3}$ REST2 MD simulation. Contacts between two residues are defined using a distance cutoff of 12Å between Cα atoms.

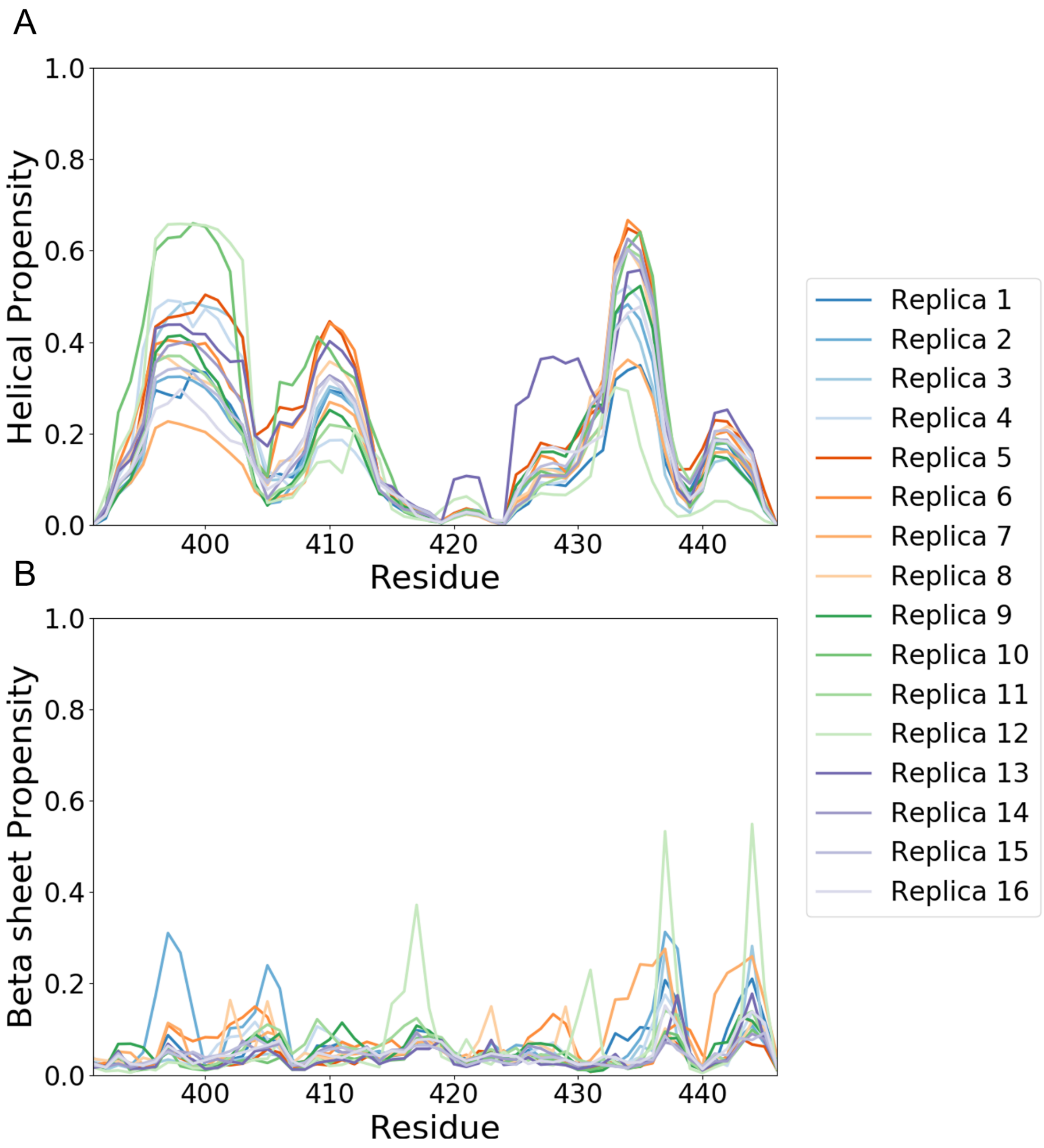


**Figure S4.** Comparison of α-helical and β-sheet propensities observed in the 16 independent demultiplexed replicas of an apo Tau-5$_{R2_R3}$ REST2 MD simulation. Secondary structure content is calculated by the DSSP algorithm. Replicas 6, 11, and 12 show the largest deviations in secondary structure propensity from the average helical propensity across all demultiplexed replicas, but still contain helical propensities in the same regions, differing only in relative populations

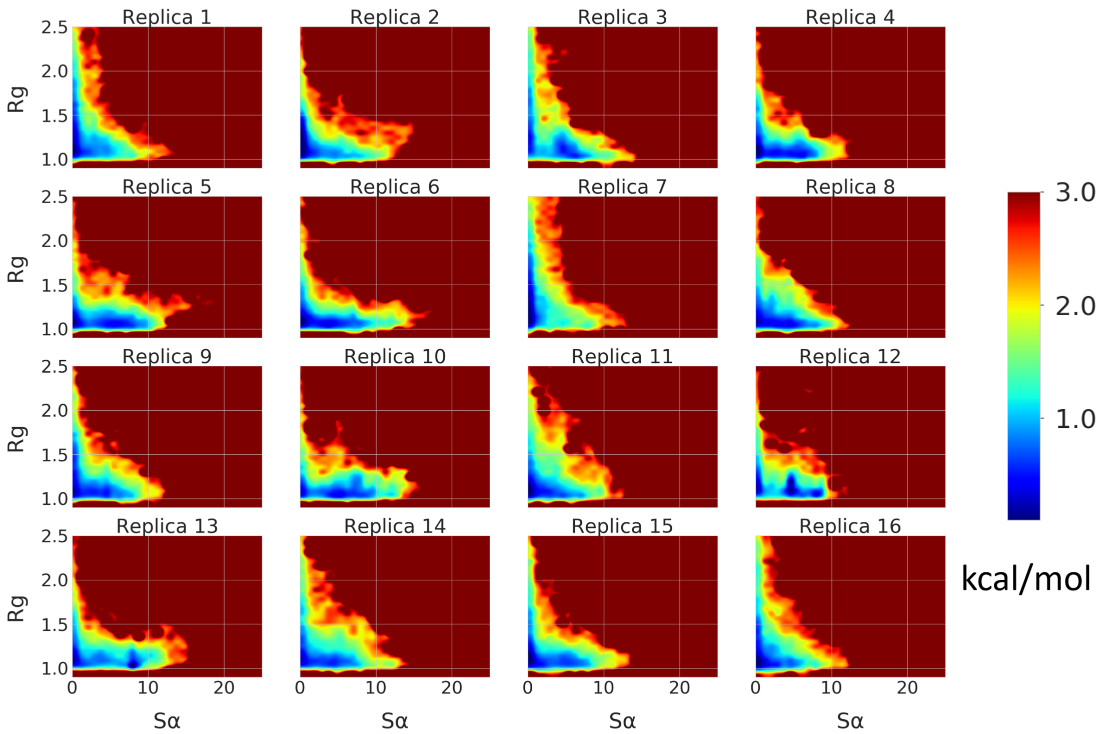


**Figure S5**. Comparison of free energy surfaces of Tau-$5_{R2\_R3}$ conformations as a function of the α-helical order parameter Sα and radius of gyration ($R_g$) for the 16 demultiplexed replicas of an apo Tau-$5_{R2\_R3}$ REST2 MD simulation. $R_g$ is reported in nm. Replica 7 samples conformations with substantially smaller Sα values than the other demultiplexed replicas.

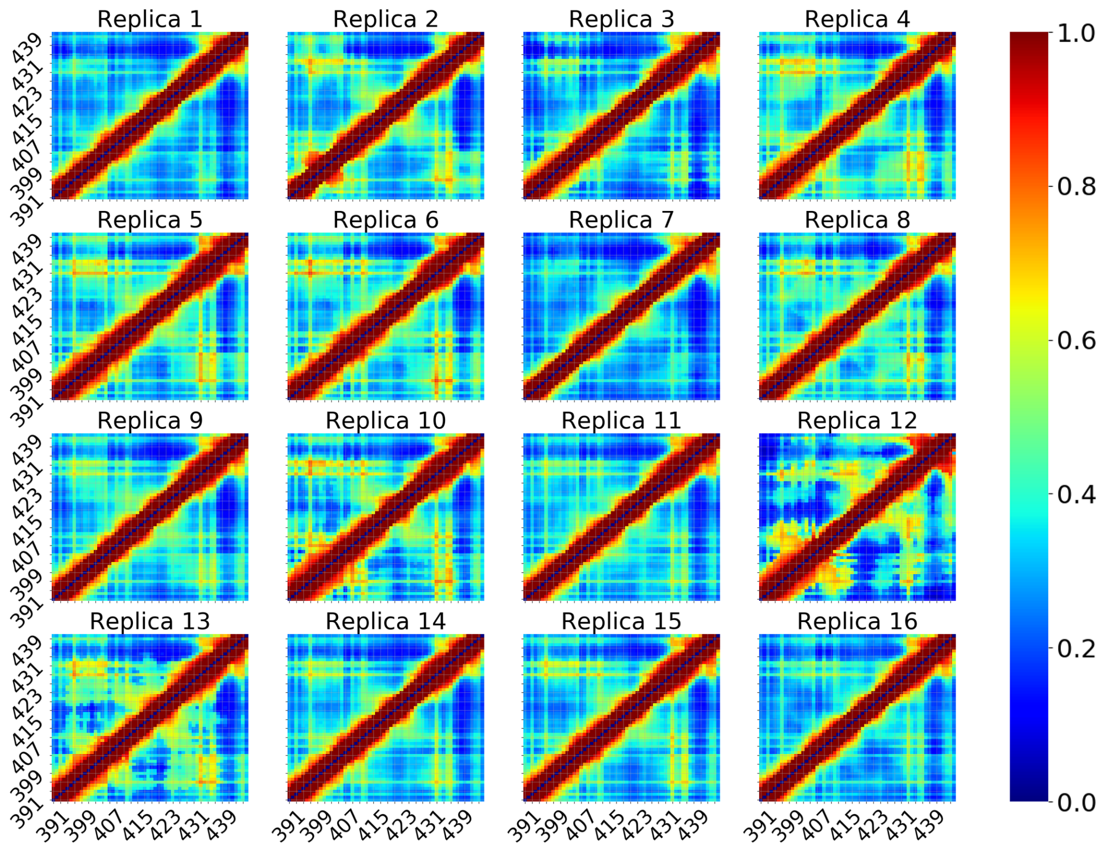


**Figure S6**. Comparison of the intramolecular contact probabilities observed in the 16 demultiplexed replicas of an apo Tau-5$_{R2_R3}$ REST2 MD simulation. Contacts between two residues were defined using a distance cutoff of 12Å between Cα atoms. Replica 7 shows substantially less populated contacts the R2 and R3 regions relative to the other replicas, and Replica 12 shows an elevated contact propensity between residues 390-395 and residues 400-405 within the R2 region.

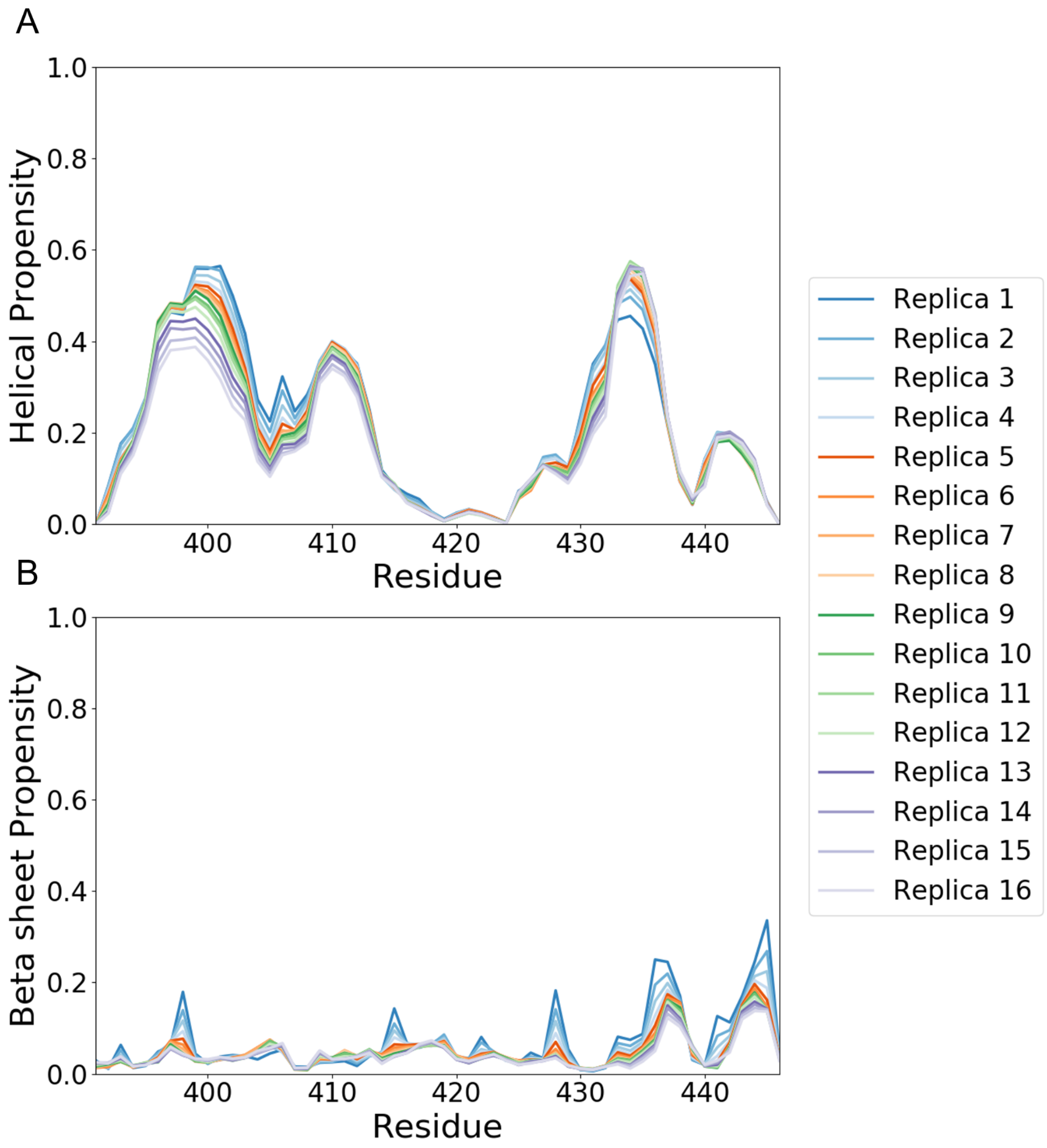


**Figure S7.** Comparison of α-helical and β-sheet propensities of Tau-$5_{R2\_R3}$ observed in the 16 solute temperature rungs of a REST2 MD simulations of Tau-$5_{R2\_R3}$ in the presence of EPI-002. Secondary structure content is calculated by the DSSP algorithm.

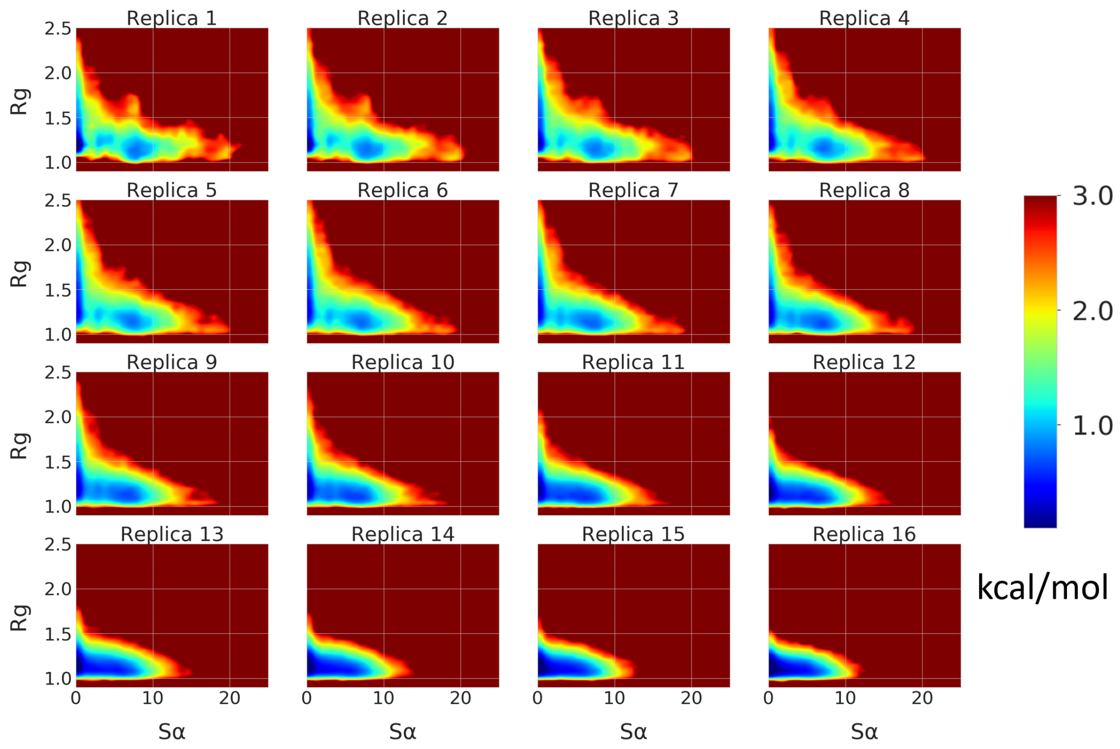


**Figure S8**. Comparison of free energy surfaces of Tau-$5_{R2\_R3}$ conformations as a function of the α-helical order parameter Sα and radius of gyration ($R_g$) for the 16 solute temperature rungs of a REST2 MD simulation of Tau-$5_{R2\_R3}$ in the presence of EPI-002. $R_g$ is reported in nm.

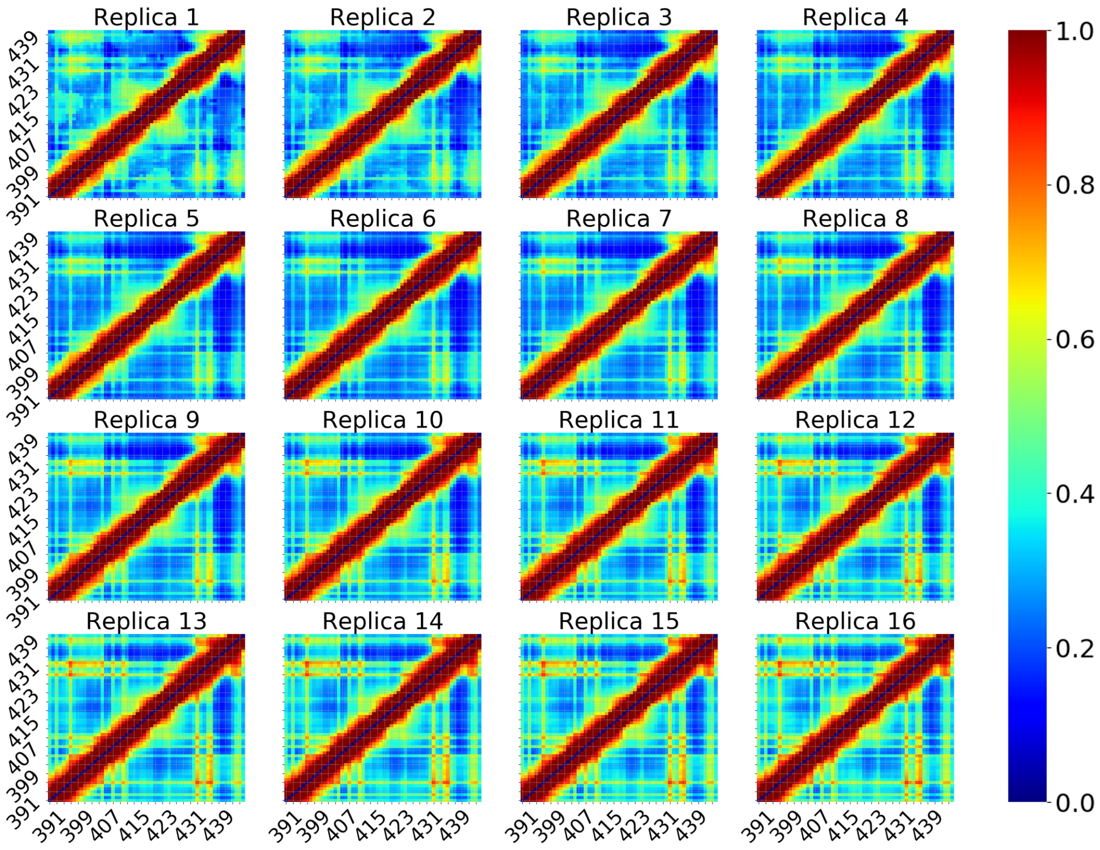


**Figure S9**. Comparison of the intramolecular contact probabilities observed in the 16 solute temperature rungs of a REST2 MD simulation of Tau-$5_{R2\_R3}$ in the presence of EPI-002. Contacts between two residues were defined using a distance cutoff of 12Å between Cα atoms.

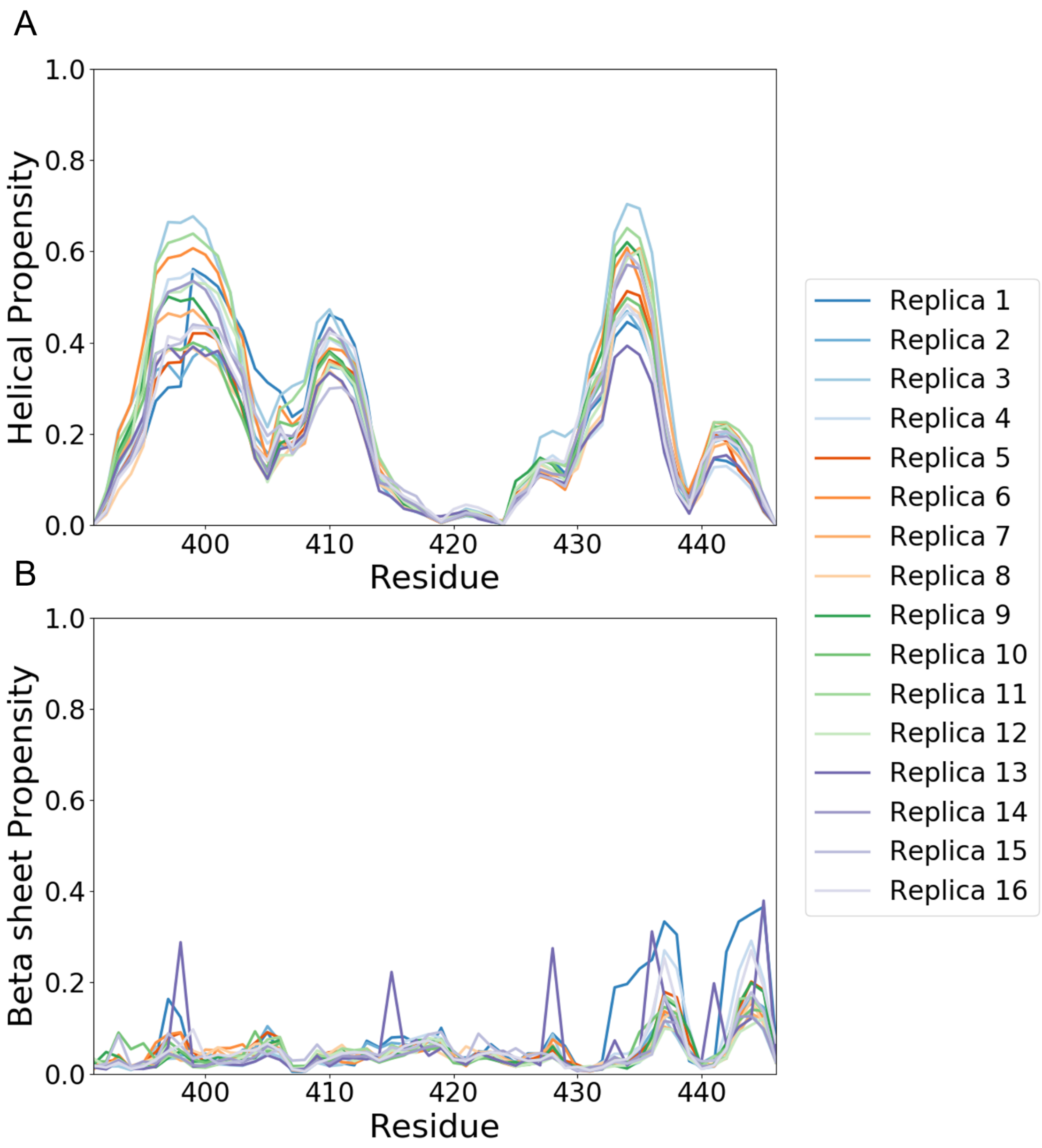


**Figure S10.** Comparison of α-helical and β-sheet propensities of Tau-$5_{R2\_R3}$ observed in the 16 demultiplexed replicas of a REST2 MD simulations of Tau-$5_{R2\_R3}$ in the presence of EPI-002. Secondary structure content is calculated by the DSSP algorithm.

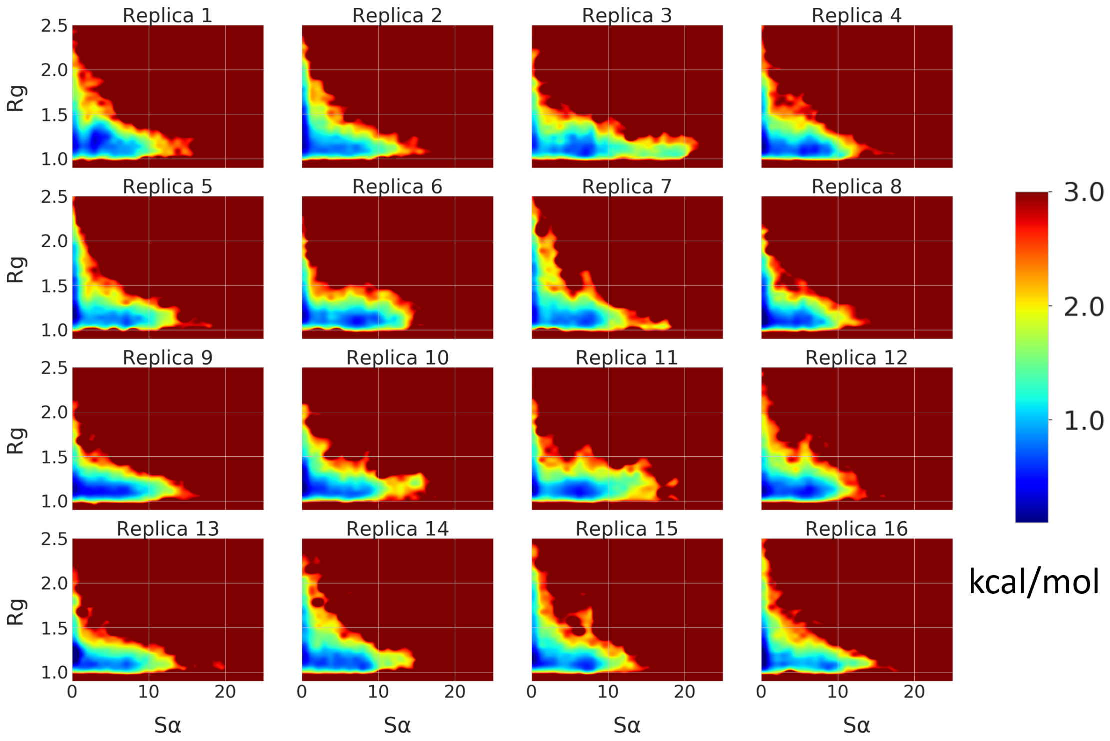


**Figure S11**. Comparison of free energy surfaces of Tau-$5_{R2\_R3}$ conformations as a function of the α-helical order parameter Sα and radius of gyration ($R_g$) for the 16 demultiplexed replicas of a REST2 MD simulations of Tau-$5_{R2\_R3}$ in the presence of EPI-002. $R_g$ is reported in nm.

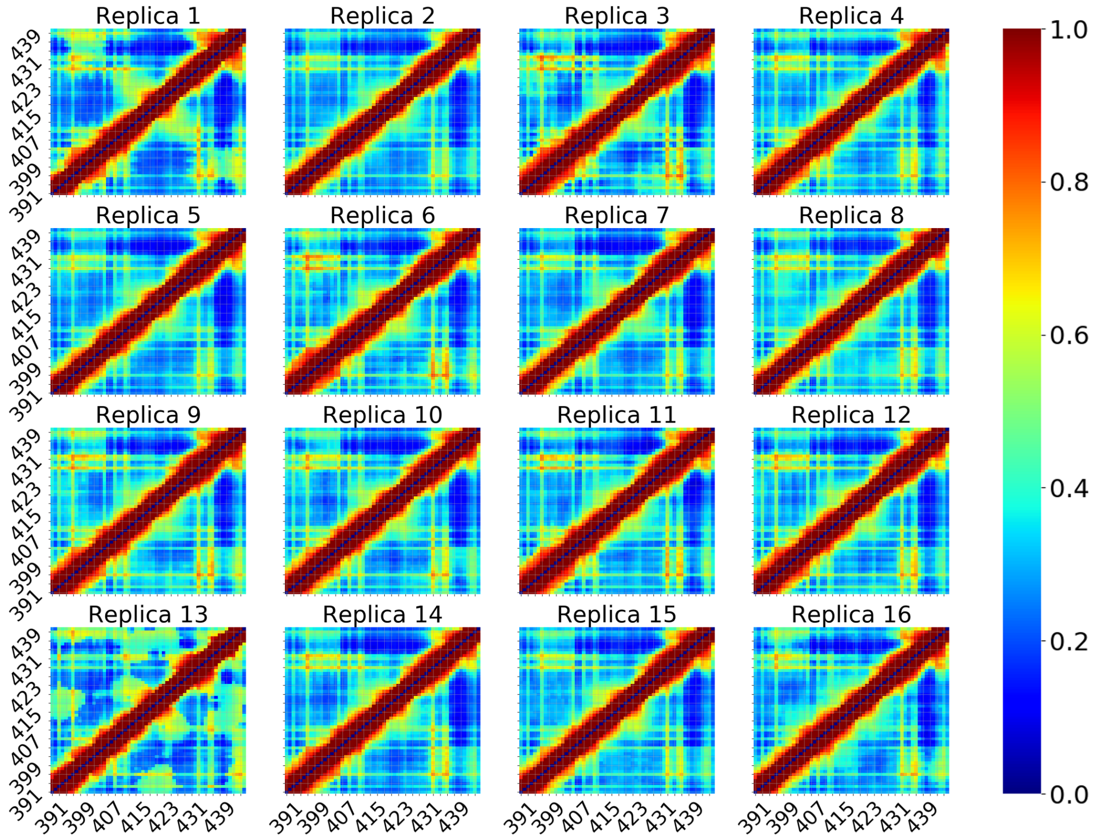


**Figure S12**. Comparison of the intramolecular contact probabilities observed in the 16 demultiplexed replicas of a REST2 MD simulations of Tau-5$_{R2_R3}$ in the presence of EPI-002. Contacts between two residues were defined using a distance cutoff of 12Å between Cα atoms.

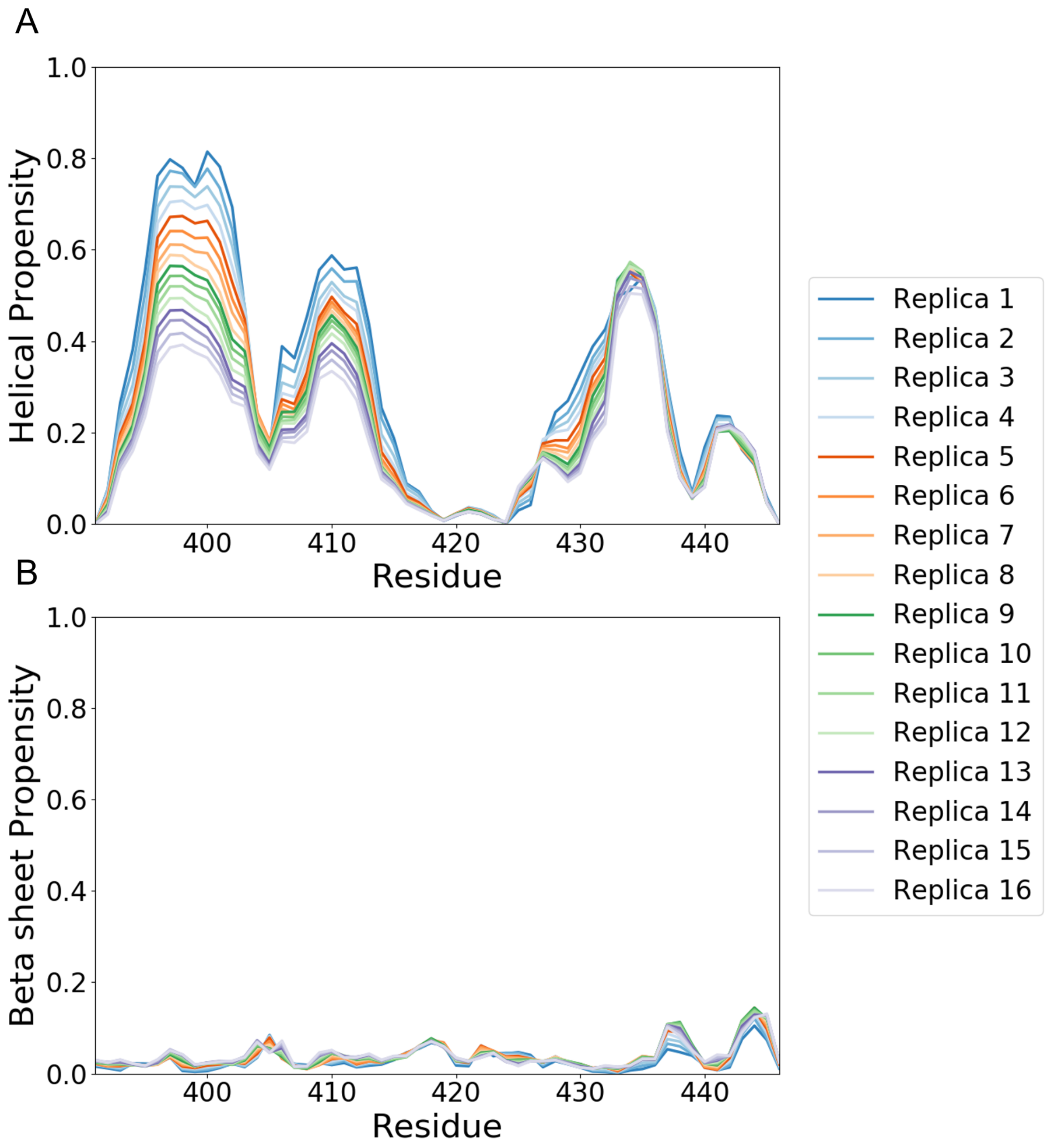


**Figure S13.** Comparison of α-helical and β-sheet propensities of Tau-$5_{R2\_R3}$ observed in the 16 solute temperature rungs of a REST2 MD simulations of Tau-$5_{R2\_R3}$ in the presence of EPI-7170. Secondary structure content is calculated by the DSSP algorithm.

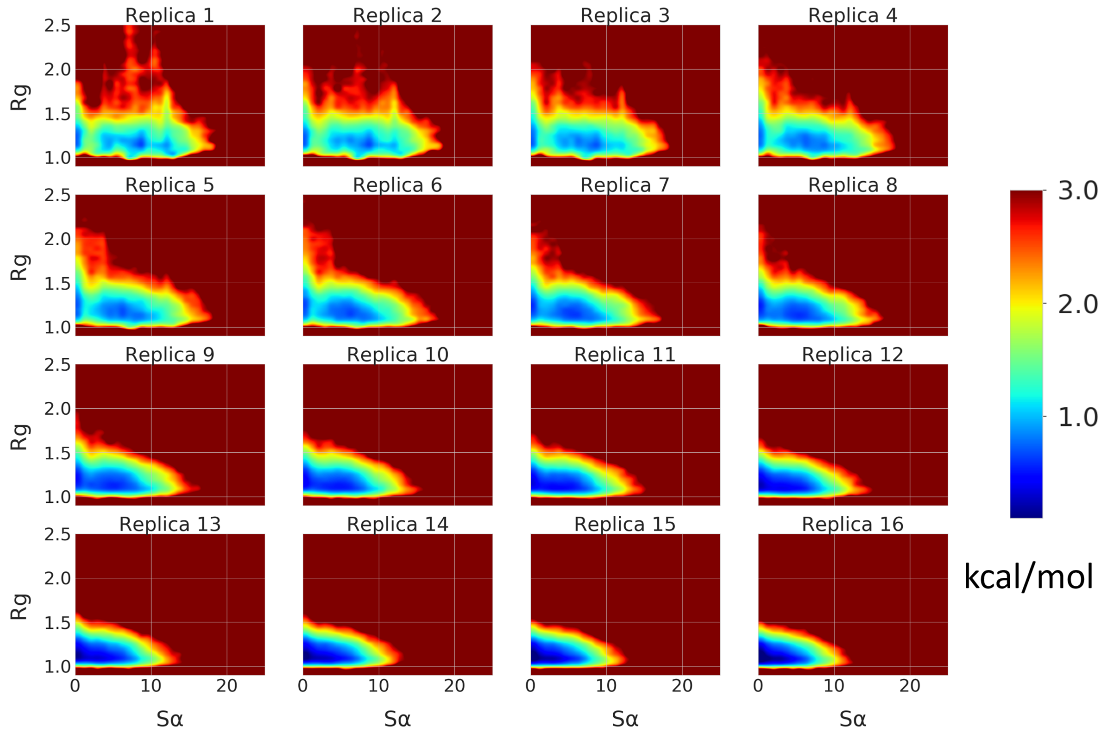


**Figure S14**. Comparison of free energy surfaces of Tau-$5_{R2\_R3}$ conformations as a function of the α-helical order parameter Sα and radius of gyration ($R_g$) for the 16 solute temperature rungs of a REST2 MD simulation of Tau-$5_{R2\_R3}$ in the presence of EPI-7170.

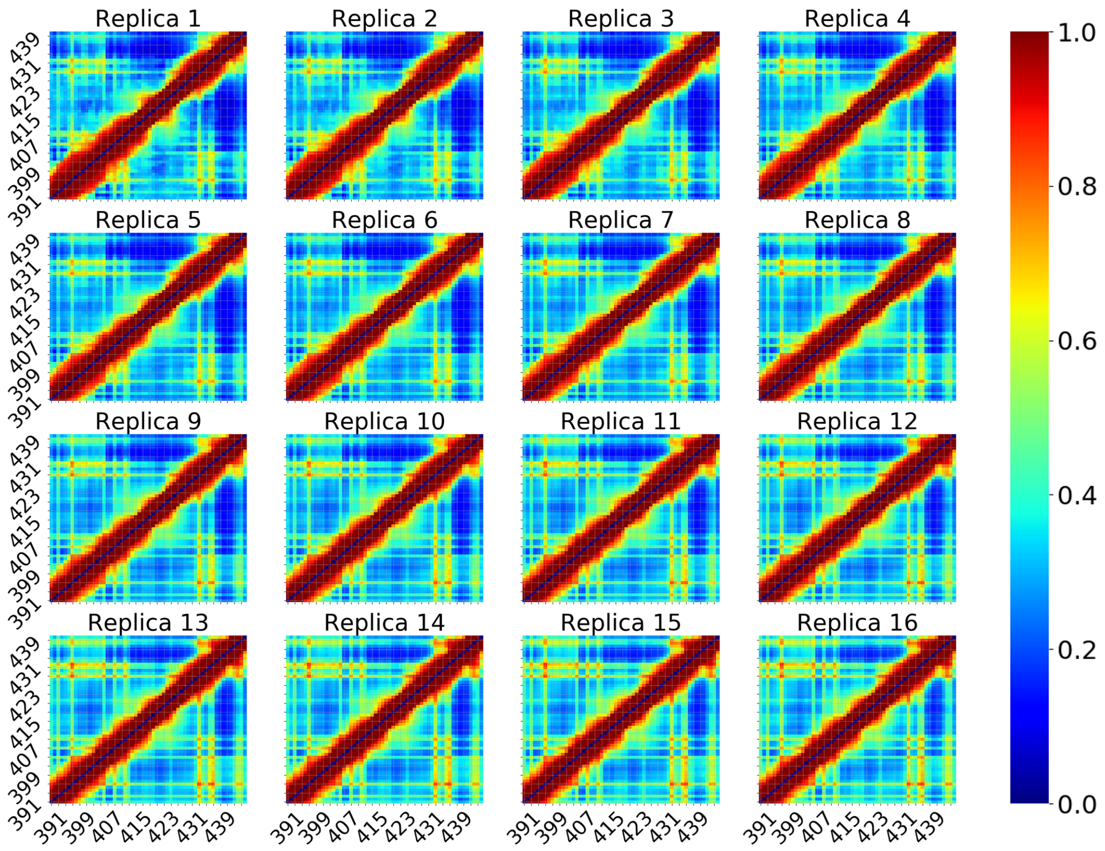


**Figure S15**. Comparison of the intramolecular contact probabilities observed in the 16 solute temperature rungs of a REST2 MD simulation of Tau-$5_{R2\_R3}$ in the presence of EPI-7170. Contacts between two residues were defined as occurring using a distance cutoff of 12Å between Cα atoms.

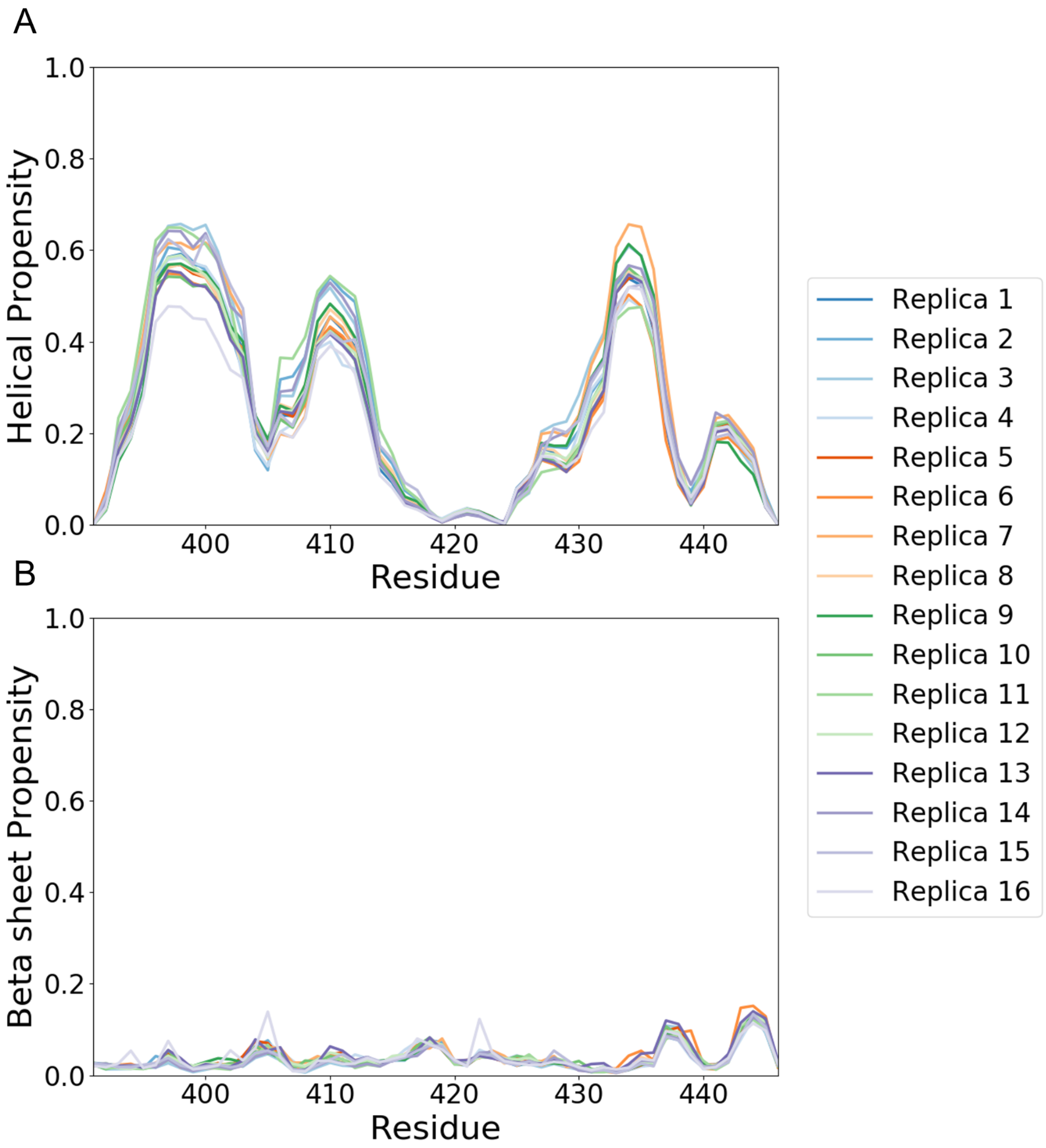


**Figure S16.** Comparison of α-helical and β-sheet propensities of Tau-$5_{R2\_R3}$ observed in the 16 demultiplexed replicas of a REST2 MD simulations of Tau-$5_{R2\_R3}$ in the presence of EPI-7170. Secondary structure content is calculated by the DSSP algorithm.

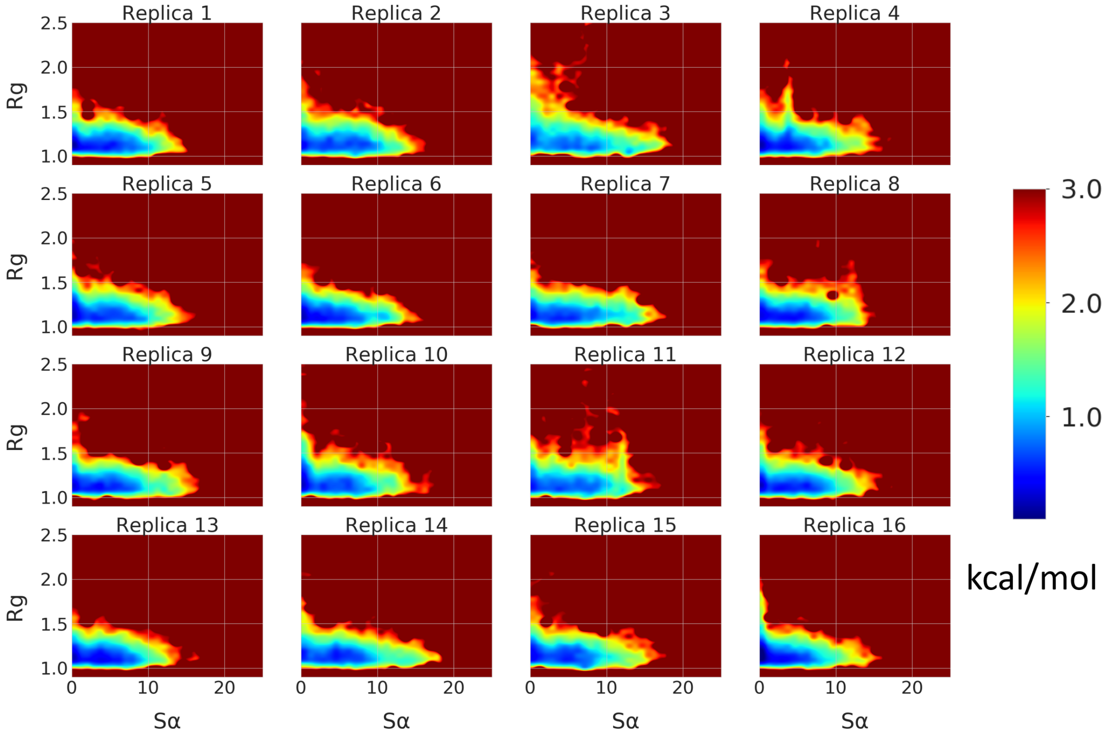


**Figure S17**. Comparison of free energy surfaces of Tau-$5_{R2\_R3}$ conformations as a function of the α-helical order parameter Sα and radius of gyration ($R_g$) for the 16 demultiplexed replicas of a REST2 MD simulations of Tau-$5_{R2\_R3}$ in the presence of EPI-7170. $R_g$ is reported in nm.

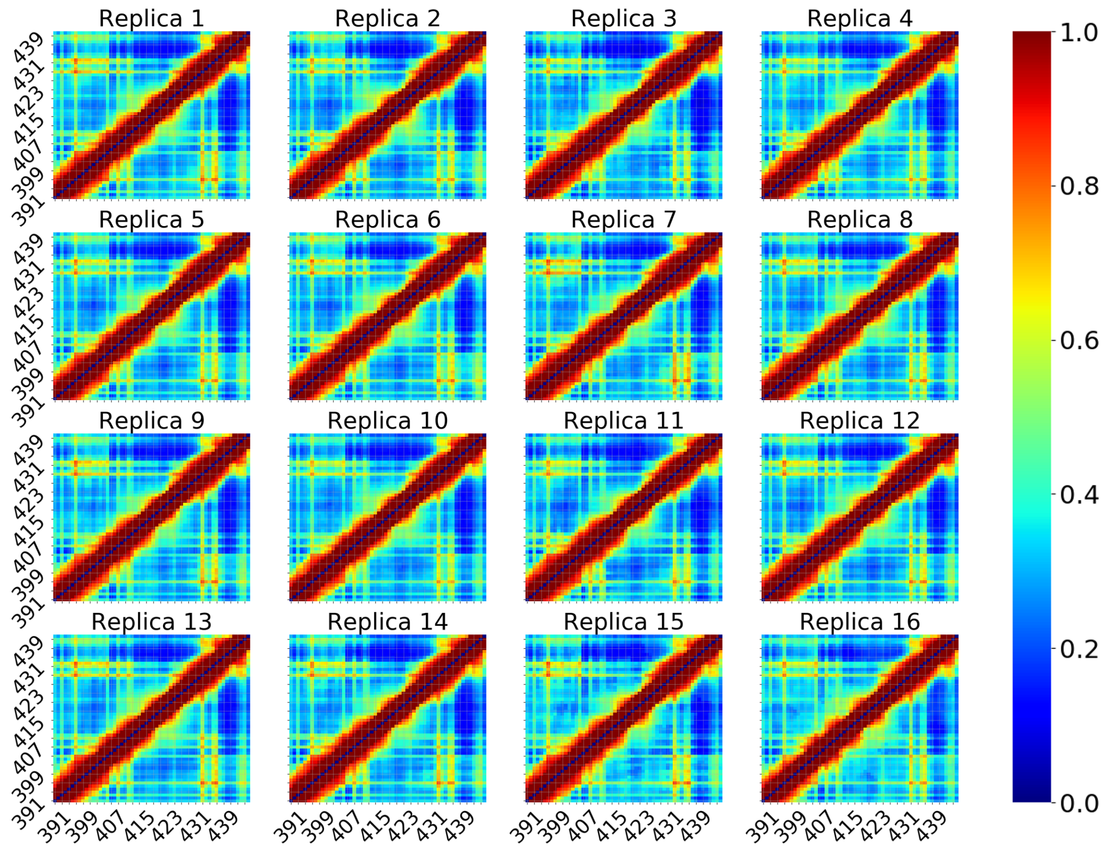


**Figure S18.** Comparison of the intramolecular contact probabilities observed in the 16 demultiplexed replicas of a REST2 MD simulations of Tau-5$_{R2_R3}$ in the presence of EPI-7170. Contacts between two residues were defined using a distance cutoff of 12Å between Cα atoms.

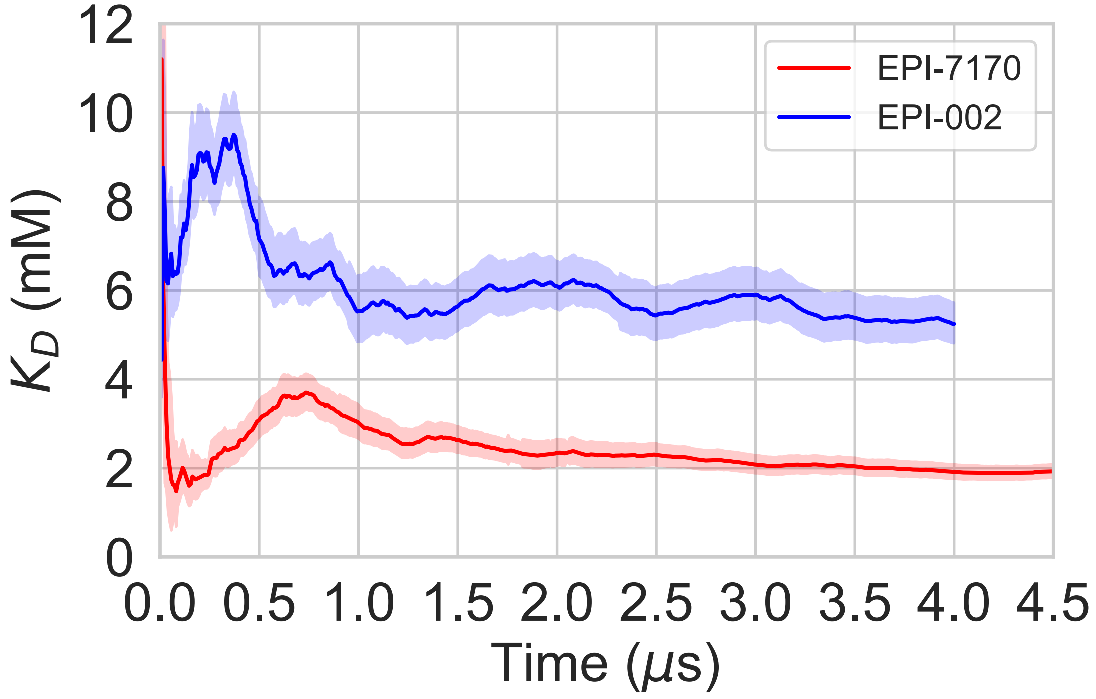


**Figure S19.** Convergence of calculated $K_D$ values for simulations of Tau-$5_{R2\_R3}$ in the presence of EPI-002 (Blue) and EPI-7170 (Red). Error estimates at each time point were calculated from a blocking analysis of the bound fraction of each compound using all frames before that time point.

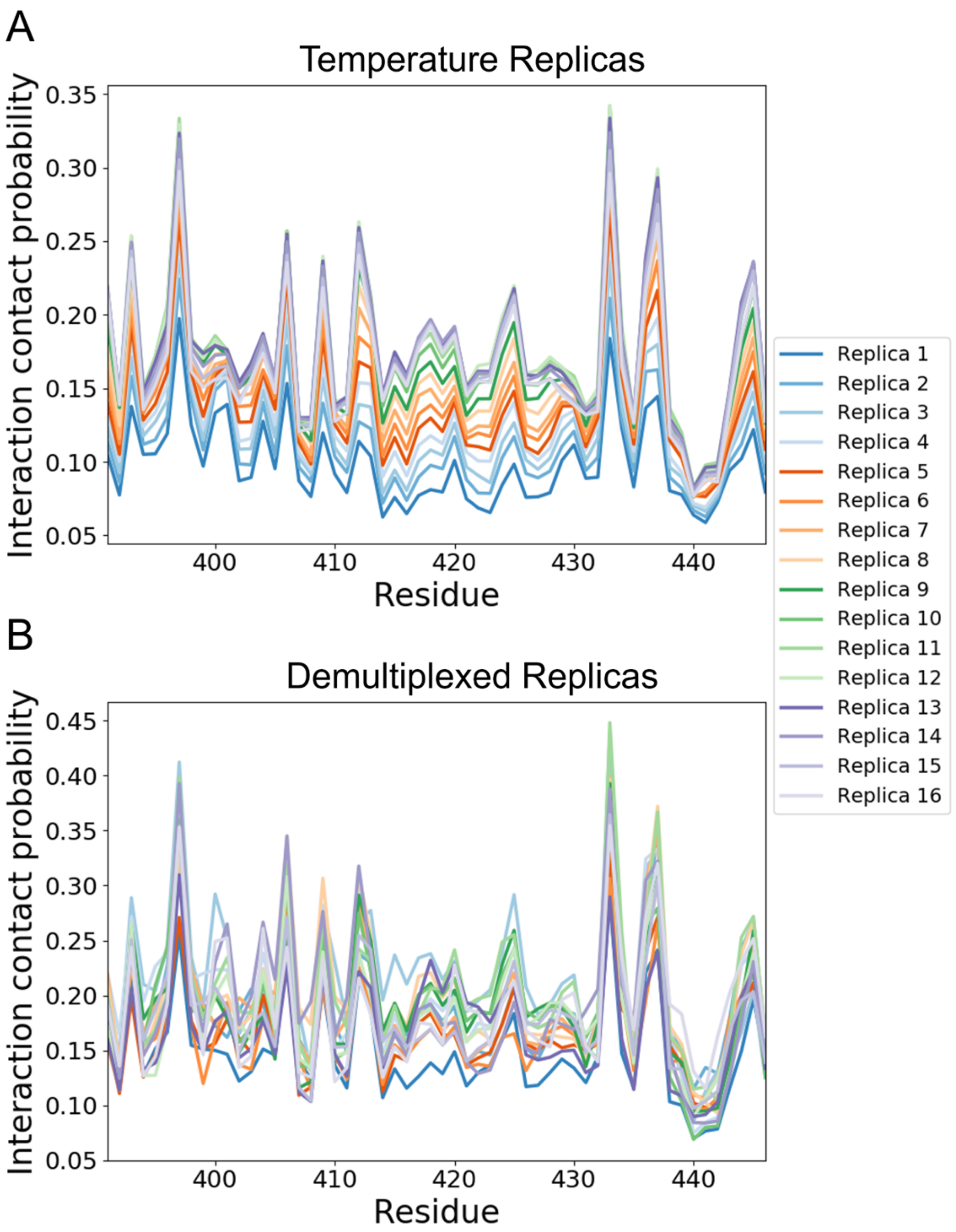


**Figure S20.** A) Ligand:protein intermolecular contact probabilities observed in the 16 solute temperature rungs from a REST2 MD simulation of Tau-$5_{R2\_R3}$ in the presence of EPI-002. Contacts between are defined between EPI-002 and Tau-$5_{R2\_R3}$ residues using a cutoff of 6Å between heavy atoms. B) Ligand:protein intermolecular contact probabilities observed in the 16 demultiplexed replicas.

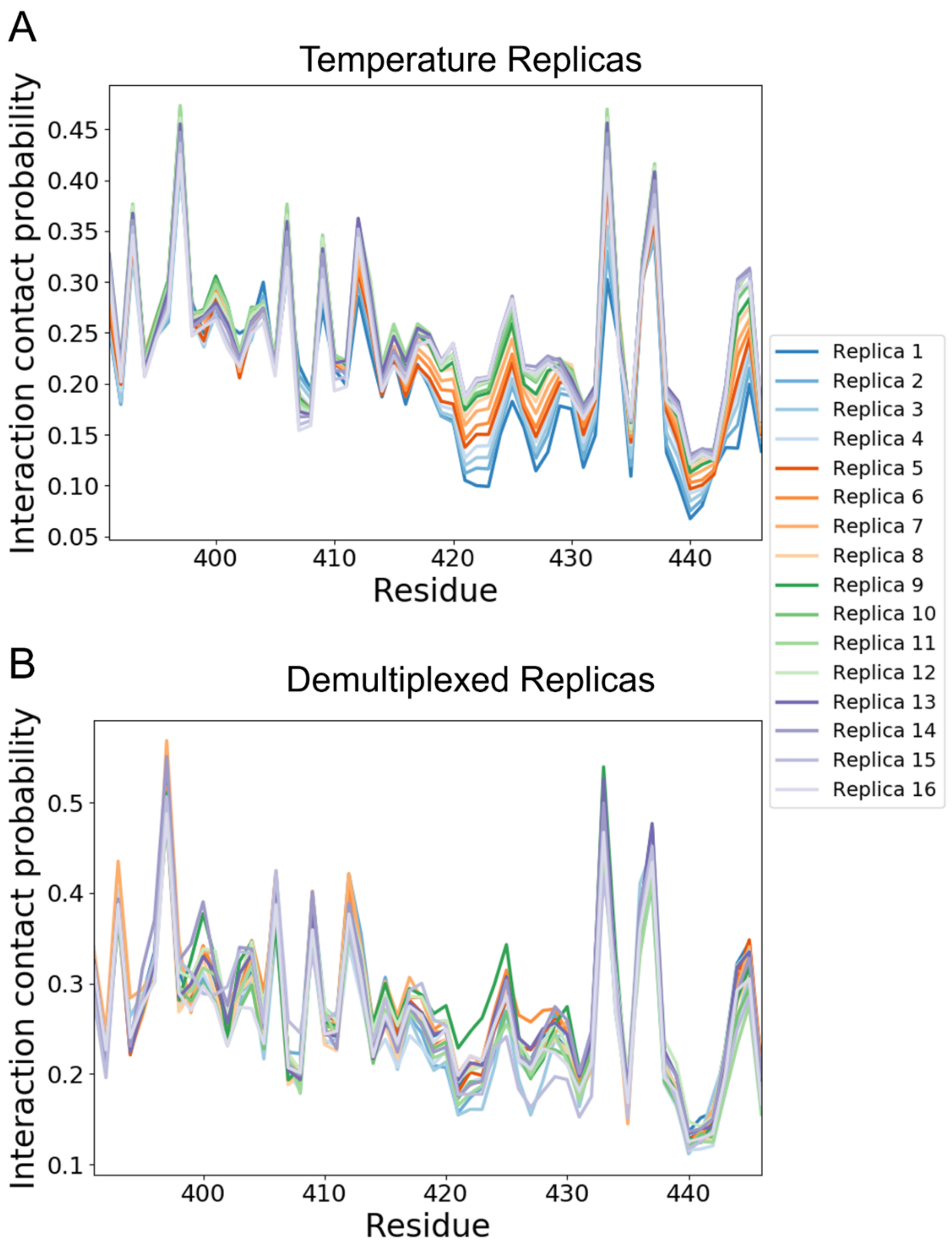


**Figure S21.** A) Ligand:protein intermolecular contact probabilities observed in the 16 solute temperature rungs from a REST2 MD simulation of Tau-$5_{R2\_R3}$ in the presence of EPI-7170. Contacts between are defined between EPI-7170 and Tau-$5_{R2\_R3}$ residues using a cutoff of 6Å between closest heavy atoms. B) Ligand:protein intermolecular contact probabilities observed in the 16 demultiplexed replicas.

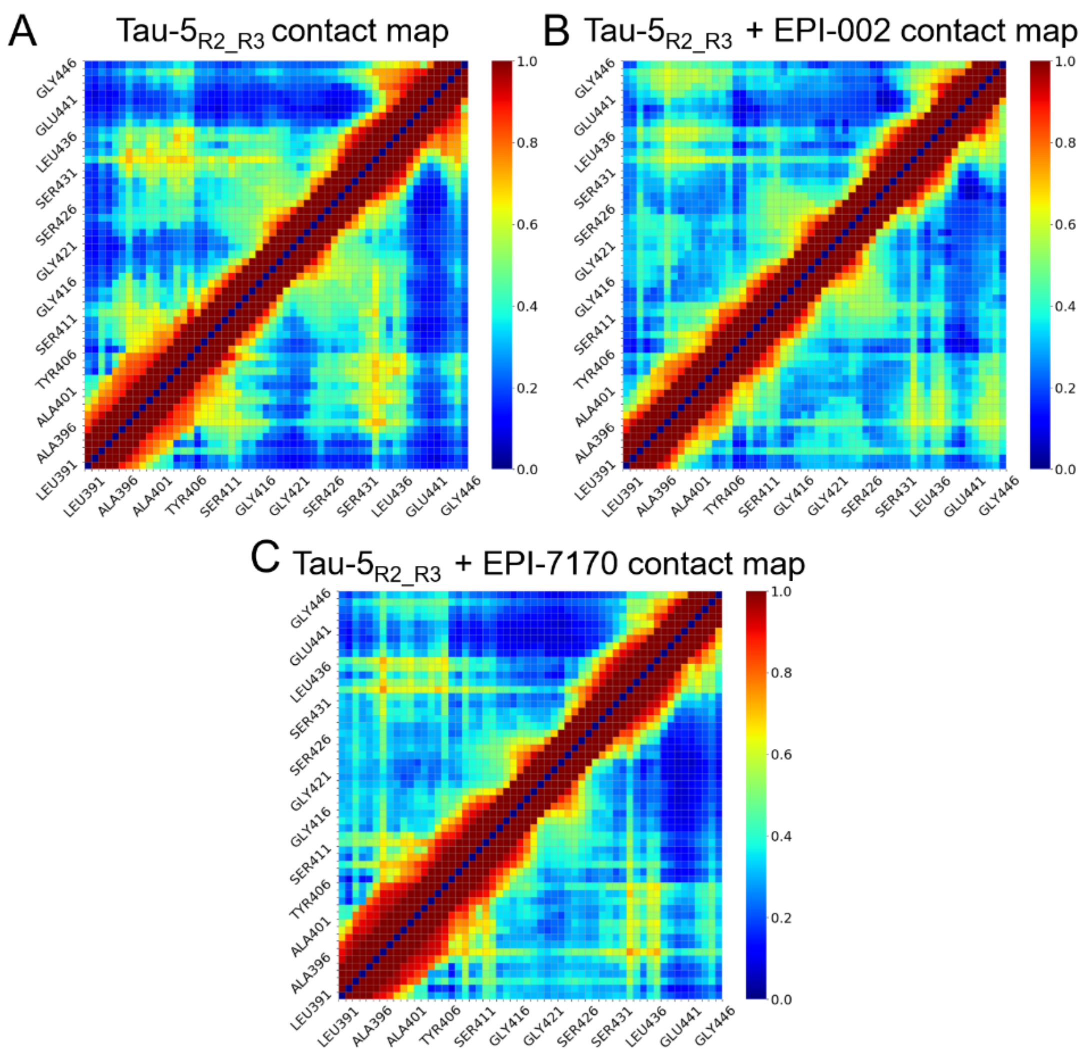


**Figure S22.** Comparison of intramolecular contact probabilities of the 300K solute temperature replicas of REST2 MD simulations of apo Tau-5$_{R2_R3}$, the EPI-002:Tau-5$_{R2_R3}$ bound ensemble and the EPI-7170:Tau-5$_{R2_R3}$ bound ensemble. Contacts between residues are defined using a cutoff distance of 12Å between Cα atoms.

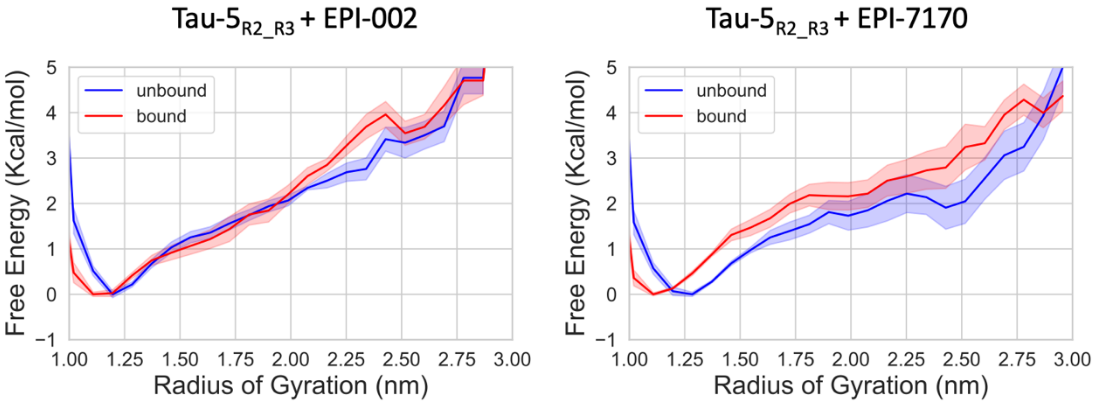


**Figure S23.** Comparison of the Radius of Gyration ($R_g$) in bound and unbound states from REST2 MD simulations of Tau-$5_{R2\_R3}$ in the presence of EPI-002 and Tau-$5_{R2\_R3}$ in the presence of EPI-7170. Bound states are defined as all frames containing at least 1 intermolecular contact (heavy atom pairs within 6Å) between a ligand and Tau-$5_{R2\_R3}$.

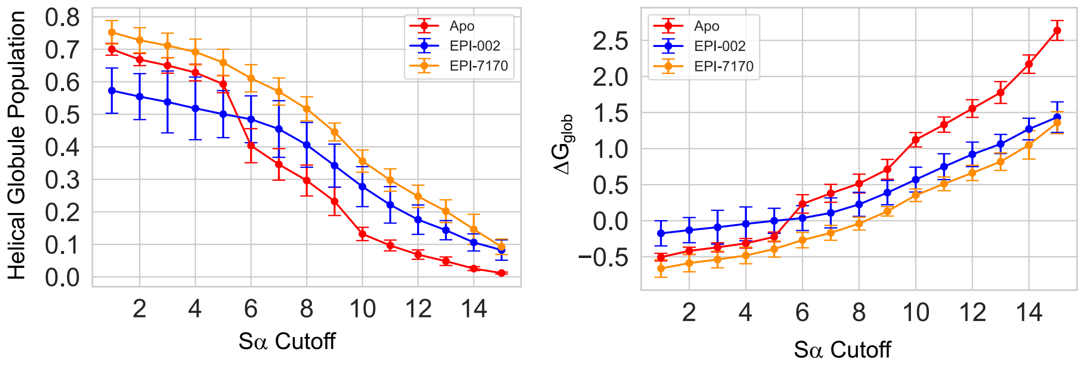


**Figure S24:** Helical globule population and stability ($\Delta G_{glob}$) relative to non-globule states in MD simulations of Tau-$5_{R2\_R3}$ in its apo form and bound to EPI-002 and EPI-7170 as a function of the Sα cutoff used in the definition of the helical globule state. Helical globule states are defined as all conformations with $R_g$<1.3nm and Sα values greater than Sα cutoff.

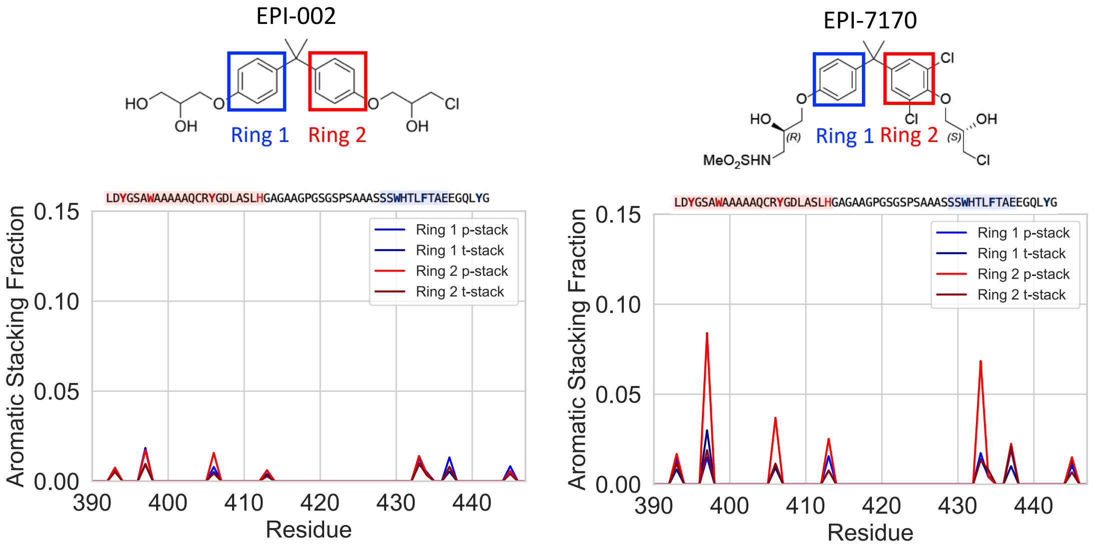


**Figure S25.** Comparison of parallel stacking (p-stack) and t-stacking (t-stack) populations in the EPI-002:Tau-5$_{R2_R3}$ and EPI-7170:Tau-5$_{R2_R3}$ bound ensembles. The dichlorinated phenyl ring of EPI-7170 has a substantially larger population of parallel stacked conformations than the corresponding non-chlorinated phenyl ring in EPI-002.

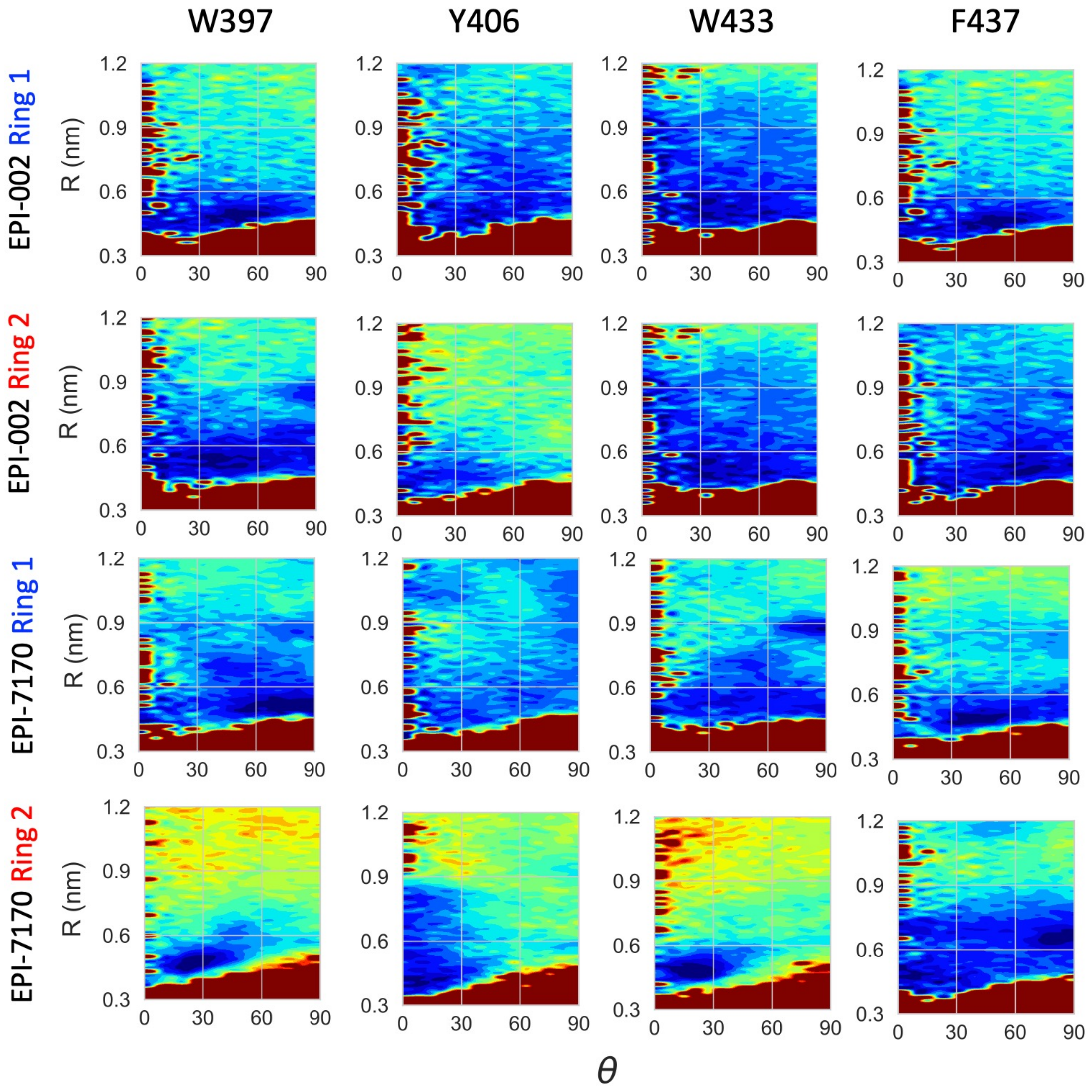


**Figure S26.** Comparison of orientations of aromatic interactions in the EPI-002:Tau-$5_{R2\_R3}$ and EPI-7170:Tau-$5_{R2\_R3}$ bound ensembles. The stacking coordinate system is described in the methods section and Fig. 5 of the main text. Ring definitions are reported in Figure S25. Free energy surfaces are shown as a function of the distance R between ring centers and the angle θ formed between the ring normal vectors

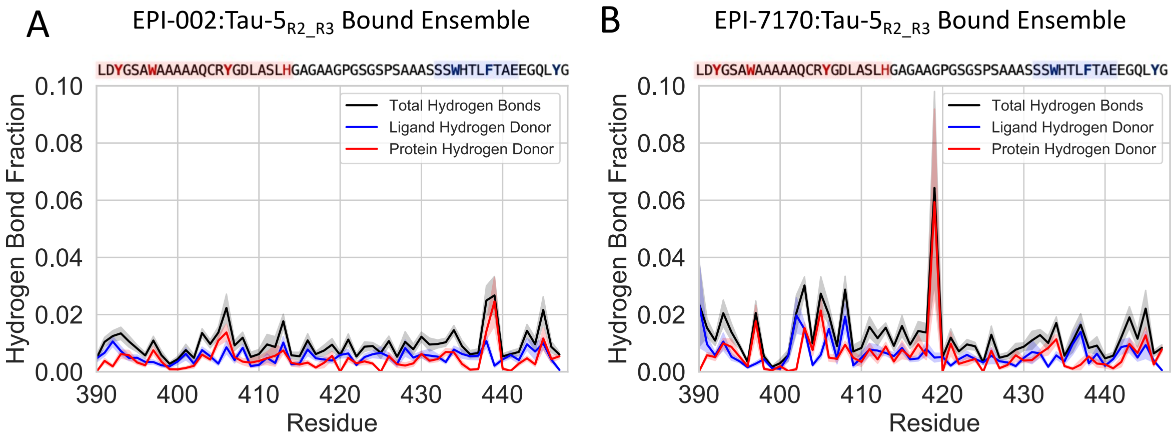


**Figure S27.** Comparison of intermolecular hydrogen bond populations observed in the EPI-002:Tau-5$_{R2_R3}$ and EPI-7170:Tau-5$_{R2_R3}$ bound ensembles. Shaded regions indicate statistical errors estimates from a blocking analysis. The total fraction of hydrogen bonds for each residue are shown in black, hydrogen bonds with ligand atoms serving as hydrogen donors are shown in blue and hydrogen bonds with protein atoms serving as hydrogen donors are shown in red. We note the most populated hydrogen bond in the EPI-7170:Tau-5$_{R2_R3}$ bound ensemble (the hydrogen bond between the backbone amide of G419 and the oxygen atom in the chlorohydrin group of EPI-7170) has a large statistical error estimate (6.4$\pm$3.4%) as it is predominantly populated in a contiguous 900ns portion of the REST2 MD trajectory in a relatively narrow subset of bound conformations.
